# Traits of bathy phytochromes and application to bacterial optogenetics

**DOI:** 10.1101/2025.05.05.652137

**Authors:** Cornelia Böhm, Kimmo Lehtinen, Elina Multamäki, Roosa Vanhatalo, Oscar Brander, Stefanie S. M. Meier, Jessica Rumfeldt, Andreas Möglich, Heikki Takala

## Abstract

Phytochromes are photoreceptors sensitive to red and far-red light found in a wide variety of organisms, including plants, fungi, and bacteria. Bacteriophytochromes (BphPs) can be switched between a red light-sensitive Pr state and a far-red light-sensitive Pfr state by illumination. In so-called prototypical BphPs, the Pr state functions as the thermally favoured resting state, whereas Pfr is more stable in bathy BphPs. The prototypical *Dr*BphP from *Deinococcus radiodurans* has been shown to be compatible with different output module types. Even though red light regulated optogenetic tools are available, like the pREDusk system based on the *Dr*BphP photosensory module, far-red light-modulated variants are still rare. Here, we study the underlying contributors to bathy over prototypical BphP behaviour by way of various chimeric constructs between pREDusk and representative bathy BphPs. We pinpoint shared traits of the otherwise heterogenous subgroup of bathy BphPs, and highlight the importance of the sensor-effector linker in light modulation of histidine kinase activity. Informed by these data, we introduce the far-red light-activated system “pFREDusk”, based on a histidine kinase activity governed by a bathy photosensory module. With this tool, we expand the optogenetic toolbox into wavelengths of increased sample and tissue penetration.

## INTRODUCTION

Light is an essential environmental stimulus that greatly affects most living organisms. Nature has developed a multitude of photoreceptor proteins capable of reacting to a wide range of wavelengths. In these photoreceptors, photosensory modules (PSMs) have evolved to respond to light of certain wavelengths through their specialised pigment molecules, the chromophores. Through light absorption by the chromophore and consequent photochemical and structural changes in their PSM, the photosensors translate environmental light conditions into cellular signalling events, which often include modulation of protein-protein interactions or enzymatic activity of their output module (OPM). The inherent modularity and flexibility of natural photoreceptors make them a prominent basis for the rational design of optogenetic tools, where the goal is to control specific cellular functions with light^1^. These highly modular systems enable high spatial and temporal resolution of modulation of biological processes, and are successfully used in the research of both eukaryotic and prokaryotic systems. Applications in bacteria range from control of gene expression to second messenger conversion and post-translational control^2^.

Among the family of photoreceptors, phytochromes have evolved as key sensors of red and far-red light, facilitated by bilin chromophores. Phytochrome function is known to affect numerous physiological processes in plants depending on the ratio of red to far-red light^3^ and has also been shown to be of importance in fungi^4^ and algae^5^. In bacteria, the significance of red and far-red light is highlighted by a large number of bacteriophytochrome (BphP) photoreceptors found across various phyla and in diverse environments^6,7^.

Structurally, typical BphPs are homodimers that constitute of an N-terminal PSM with a PAS (Period/ARNT/Single-minded), a GAF (cGMP phosphodiesterase/Adenylate cyclase/FhlA) and a PHY (phytochrome-specific) domain (Fig. 1A). Within the PSM, the PAS-GAF core provides the chromophore-binding pocket which is bracketed by a hairpin extension of the PHY domain, referred to as the “PHY-tongue” (PTG; Fig. 1B-C), that has been shown to play a major role in light-dependent signal transduction^8^. The chromophore itself is covalently bound to a conserved cysteine residue in the N-terminal segment (NTS), a structural element preceding the PAS domain core that is involved in the formation of a characteristic figure-of-eight knot structure^9^. Both PHY-tongue and NTS interact directly with the linear tetrapyrrole chromophore biliverdin IXα (BV), which undergoes *Z*/*E* isomerisation and concomitant rotation of its *D*-ring around its C15=C16 double bond upon illumination^10^.

**Figure 1:**
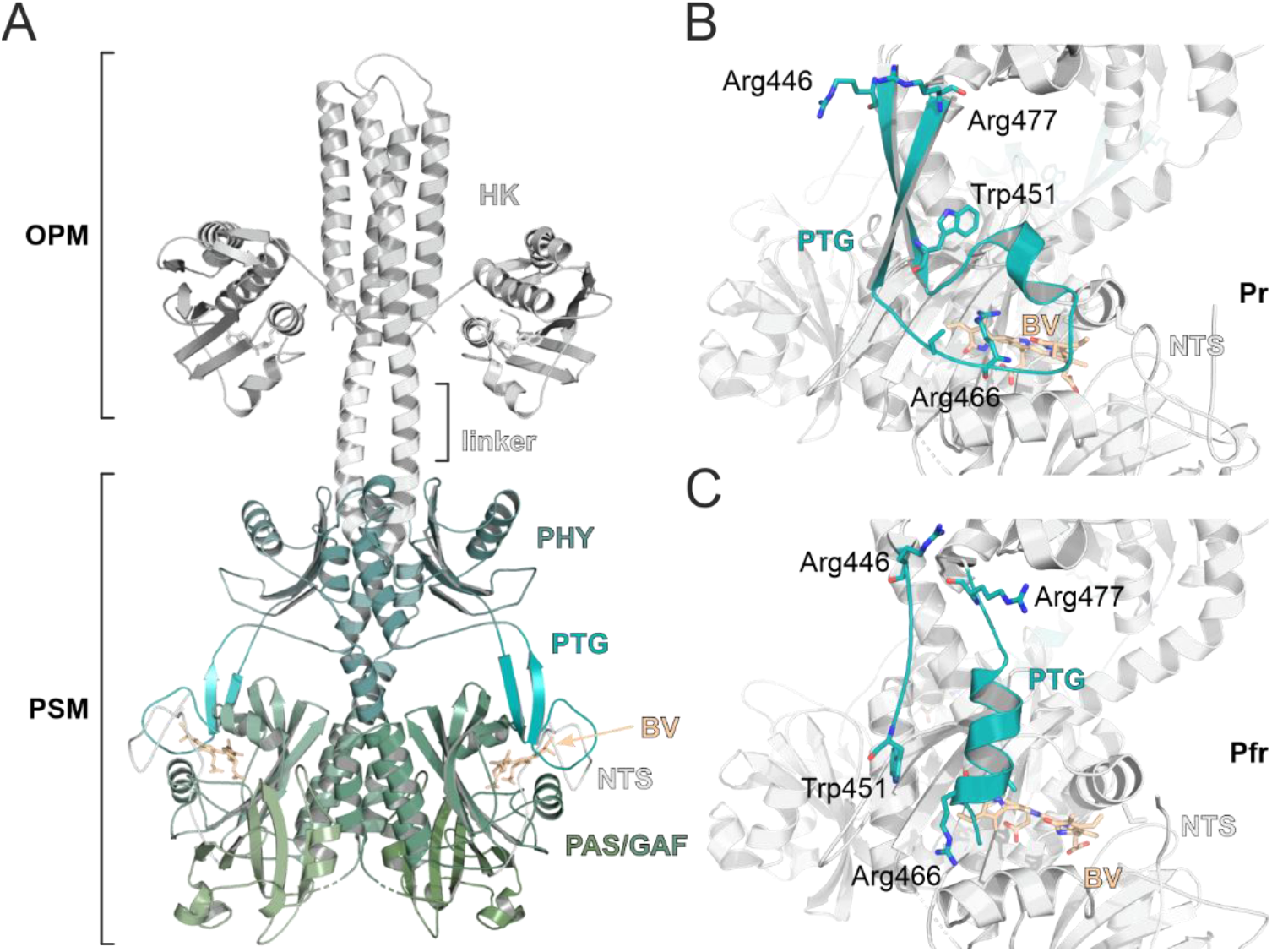
Structure of bacteriophytochrome histidine kinases. **(A)** Structural model of a bacteriophytochrome with a histidine kinase (HK) output module. The structural model is assembled from the cryo-EM structure of DrBphP in the Pr state (PDB code: 8AVW)^11^ and the HK module crystal structure of Thermotoga maritima TM0853 (PDB code: 2C2A)^12^. **(B-C)** Structure of the DrBphP PHY-tongue (PTG) in its Pr (B, PDB code: 8AVV) and Pfr state (C, PDB code: 8AVX). Selected residues and biliverdin chromophore (BV, orange) are indicated (Trp451 and Arg466 are part of the WAG and PRXSF motifs, respectively).

In the majority of known BphPs, the red light-absorbing Pr state (BV in *ZZZssa* conformation) acts as the energetically favoured resting state, which can be switched to the far-red light-absorbing Pfr state (BV in *ZZEssa*) with red light (∼650 nm). The Pfr state, on the other hand, is sensitive to far-red light (∼750 nm) and can return to Pr either through far-red light exposure or through thermal relaxation. However, BphPs are not exclusively confined to a Pr resting state: As reported by Karniol *et al*. already in 2003, some BphPs feature Pfr as a resting state and have been since termed “bathy”^13^, as opposed to “prototypical” ones. The determinants of bathy over prototypical behaviour in BphPs remain unelucidated to date, even though several bathy representatives have been identified and characterised^13–20^.

Upon BphP photoactivation, isomerisation of BV is accompanied by local structural rearrangements starting from the chromophore binding pocket, which include refolding of the PHY-tongue between β-hairpin (Pr state) to α-helix (Pfr state) conformations^21,22^. These structural changes in the PSM are further relayed to the C-terminal OPM through larger-scale rearrangements. The molecular mechanism of this signal processing depends on the individual OPM type^23,24^, which varies among BphPs but is often a histidine kinase (HK) module^7^. Typically, the HK modules of BphPs are either members of the HisKA (Interpro IPR003661) or the HWE-HK (Interpro IPR011102) families. As red/far-red light-sensing HKs, these BphPs are part of bacterial two-component systems (TCSs). TCSs consist of a HK and a response regulator (RR) protein component and are prevalent signalling pathways in response to environmental stimuli in bacteria^25,26^. There, autophosphorylation of the catalytic histidine residue of the HK is succeeded by phosphotransfer to a conserved aspartate within the receiver (REC) domain of a cognate RR. This aspartate phosphorylation then leads to an RR-mediated output response, often through transcriptional regulation^27^. The HK itself can function both as a kinase or phosphatase, and RR phosphorylation depends on the net balance between these opposing activities^28,29^.

With the growing availability of red light-regulated phytochrome-based optogenetic tools^30–33^, the demand for additional far-red and near infrared (NIR) wavelength sensitivities increases. Bathy BphPs constitute a promising basis for optogenetic systems as they are regulated by far-red light which falls within the so-called NIR tissue transparency window^34,35^. Therefore, the introduction of bathy alternatives into systems based on prototypical BphPs is a promising approach. One such prototypical tool is the red light-repressed pREDusk gene expression system^33^. The function of pREDusk is governed by the HK activity of the *Dr*F1 chimaera, which consists of the PSM from *Dr*BphP (a model BphP from *Deinococcus radiodurans*)^21^ and the HK module from *Bradyrhizobium japonicum* FixL (*Bj*FixL)^36^. In darkness, *Dr*F1 phosphorylates the cognate RR *Bj*FixJ^36^, subsequently inducing target gene expression. Red light renders *Dr*F1 net phosphatase-active, which leads to inactivation of gene expression.

Here, we study the determinants underlying bathy behaviour of BphPs with the goal to make the pREDusk tool far-red light-activatable. To design such a system, we create chimaeras between *Dr*F1 and three well-known bathy BphPs – *Agrobacterium fabrum* Agp2^13^, *Pseudomonas aeruginosa Pa*BphP^15^ and *Rhodopseudomonas palustris Rp*BphP1^19^. Whereas PHY-tongue and NTS replacements yield chimeras with prototypical characteristics, exchanging the entire *Dr*PSM for the corresponding module of bathy origin turns the entire protein component bathy. In addition to gaining further insight into the relationship between bathy and prototypical BphPs and the effects of the sensor-effector linker region, we were able to create the far-red light-activated pFREDusk tool with very low background activity in the dark and several hundred-fold upon upregulation of target gene expression with far-red light.

## RESULTS AND DISCUSSION

### Bathy BphPs are more closely conserved than BphPs in general

With the aim to generate far-red light-activatable pREDusk tool governed by a bathy version of *Dr*F1, we searched the existing literature for wild-type BphPs with published spectral properties, and found 41 entries, 11 of which are bathy. Although this implies that 25–30% of BphPs in general could be bathy, it is impossible to estimate whether this reflects all naturally occurring BphPs as the total number of known bathy BphPs remains relatively low. Moreover, many known bathy BphPs were presumably identified based on sequence similarity to the first published representatives. Nevertheless, we screened the known BphPs with bathy characteristics for conserved features on a sequence level, with the intent to identify resting state determinants. Interestingly, bathy BphPs do not strongly cluster phylogenetically among BphPs with known spectral properties (Fig. S1). When probed for differences between known bathy and prototypical BphPs, however, several features stand out as more strongly conserved within bathy BphPs than among BphPs in general.

To begin with, a characteristic that varies notably across BphP sequences is the length of the PHY-tongue. Here, the PHY-tongue is defined to extend from 5 residues preceding the conserved WAG motif (WAG–5) to 8 residues after the PRXSF motif (PRXSF+8), which correspond to residues R446 and R477 in *Dr*BphP, respectively (Fig. 1B-C). Among a total of 404 BphP sequences with a PAS-GAF-PHY architecture retrieved from InterPro, including all representatives with published spectral properties we could find, PHY-tongue lengths range from +0 to +10 residues relative to *Dr*BphP. Interestingly, the PHY-tongue elements of all known bathy BphPs are +5 residues longer than in *Dr*BphP, a length that also contains the general majority of BphPs (Fig. 2, Fig. S2A). As already noted by Xu *et al*., these variations in length are largely confined to the area preceding the PRXSF motif^37^.

**Figure 2:**
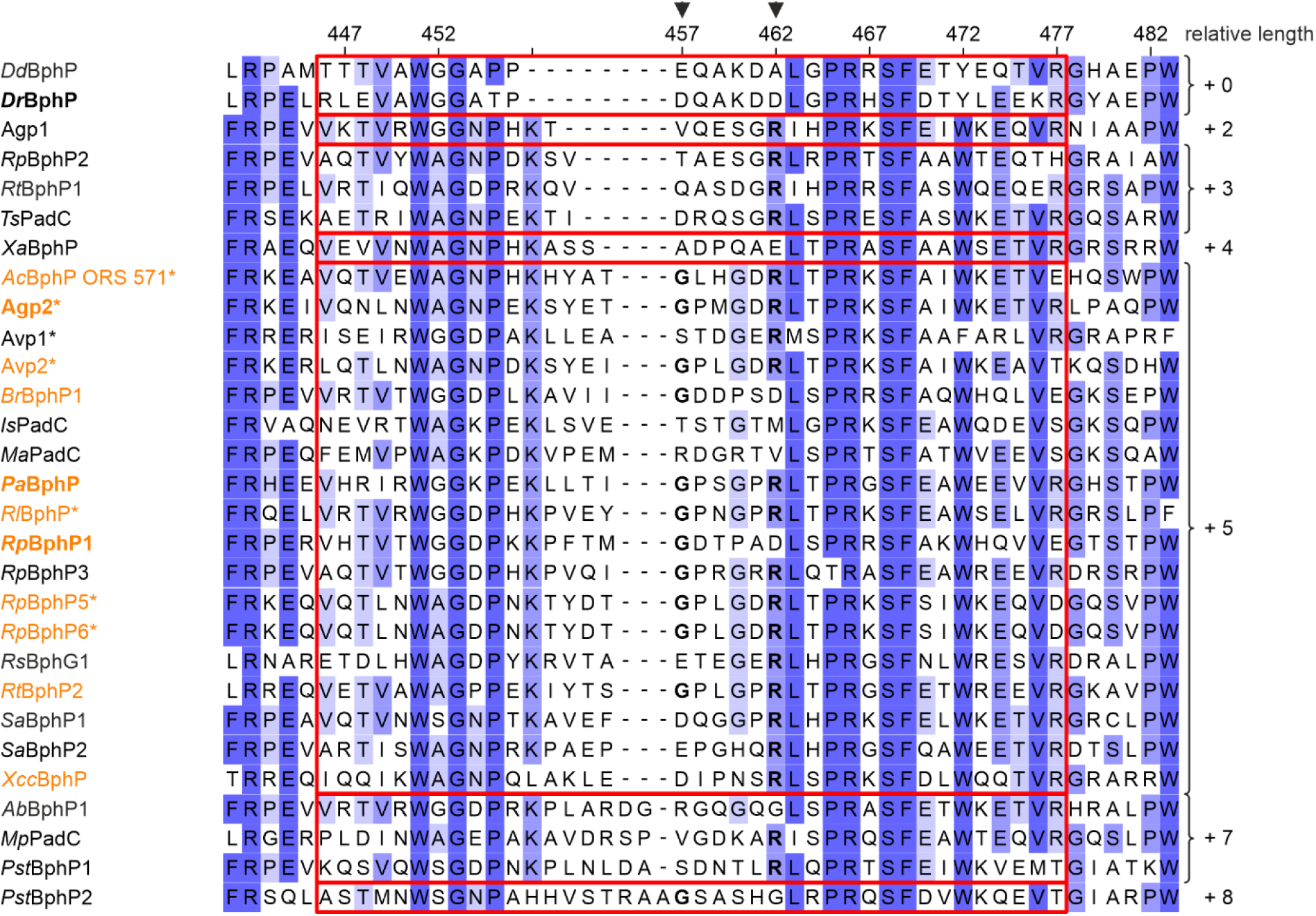
Features conserved more strictly in bathy phytochromes than in BphPs in general. Comparison of PHY-tongue lengths in representative bona fide BphPs counted from WAG–5 (DrBphP R446) to PRXSF+8 (DrBphP R477). Residues are coloured according to Jalview sequence ID (hues of blue correspond to ≥ 84%, ≥ 68%, ≥ 40%, < 40% residue conservation) based on an alignment of 404 sequences, numbering corresponds to DrBphP. HWE-HKs are marked with *, bathy BphPs in orange. PRXSF−8 glycine and PRXSF−3 arginine^38^ are marked with arrows. XccBphP, though initially classified as “bathy-like”, is also considered a bathy BphP due to the full Pfr resting state of its PSM truncation^20^. See Table S1 for a list of represented BphPs, and supplementary files for the full alignment and details on all sequences.

In addition to a previously characterised conserved arginine (PRXSF−3, D462 in *Dr*BphP)^38^, conservation of another residue within the PHY-tongue attracts further attention – a glycine at position PRXSF−8 (*Dr*BphP D457). This glycine, though not strictly conserved in known bathy BphPs (Fig. S2B), is considerably more prevalent among bathy representatives than BphPs in general (Fig. S2C). It is typically part of a loop formed at the tip of the PHY-tongue between the conserved WAG and PRXSF motifs (Fig. 1B-C), which may provide flexibility to the loop. Although the previously highlighted arginine in position PRXSF−3 (*Dr*BphP D462)^38^ and a glutamine in position PASDIP−3 (*Dr*BphP H201)^39^ are found in most bathy BphPs, not all BphPs with a glutamine or arginine in these positions are bathy. Despite a generally strongly conserved histidine in position LWGL+5 (*Dr*BphP H290) being present in all known bathy BphPs and though substitution by alanine reverts *Pa*BphP to a prototypical phytochrome^40^, no such effect is observed for a corresponding Agp2 variant^41^. On a sequence level, the BV environment in bathy BphPs therefore appears to be subtly different from prototypical BphPs, but not exclusively or distinctly so.

### PHY-tongue exchanges favour prototypical BphP behaviour

Introduction of a prototypical PHY-tongue into a *Pa*BphP-based variant has previously been reported to yield a prototypical construct^37^. Consequently, following our observations about conservation of PHY-tongue length in known bathy BphPs, we created three *Dr*F1-based chimaeras by introducing bathy PHY-tongues from Agp2, *Pa*BphP and *Rp*BphP1 into the *Dr*PSM (Fig. 3A). Interestingly, the corresponding variants *Ag*PTG, *Pa*PTG and *Rp*PTG all appear prototypical, with classical Q-band intensities and spectra resembling the parent *Dr*F1 construct (Fig. 3B-C). It is worth noting that the apparent Pfr state occupancy is affected by these PHY-tongue exchanges, with the chimaeras converting less to Pfr upon red light exposure than *Dr*F1 or wild-type *Dr*BphP^42^. This reduced Pfr-state occupancy may be partially explained by notably faster dark state recovery in *Ag*PTG, *Pa*PTG and *Rp*PTG than in *Dr*F1 (Fig. S3A,E). Another structural element of the PSM known to affect spectral properties, especially in interplay with the PHY-tongue, is the NTS^43,44^. To investigate its role in spectral behaviour, NTS exchanges were carried out for Agp2 and *Pa*BphP with and without accompanying PHY-tongue exchanges (Fig. S4A), but all resulting chimaeras appear prototypical (Fig. S4B,C).

**Figure 3:**
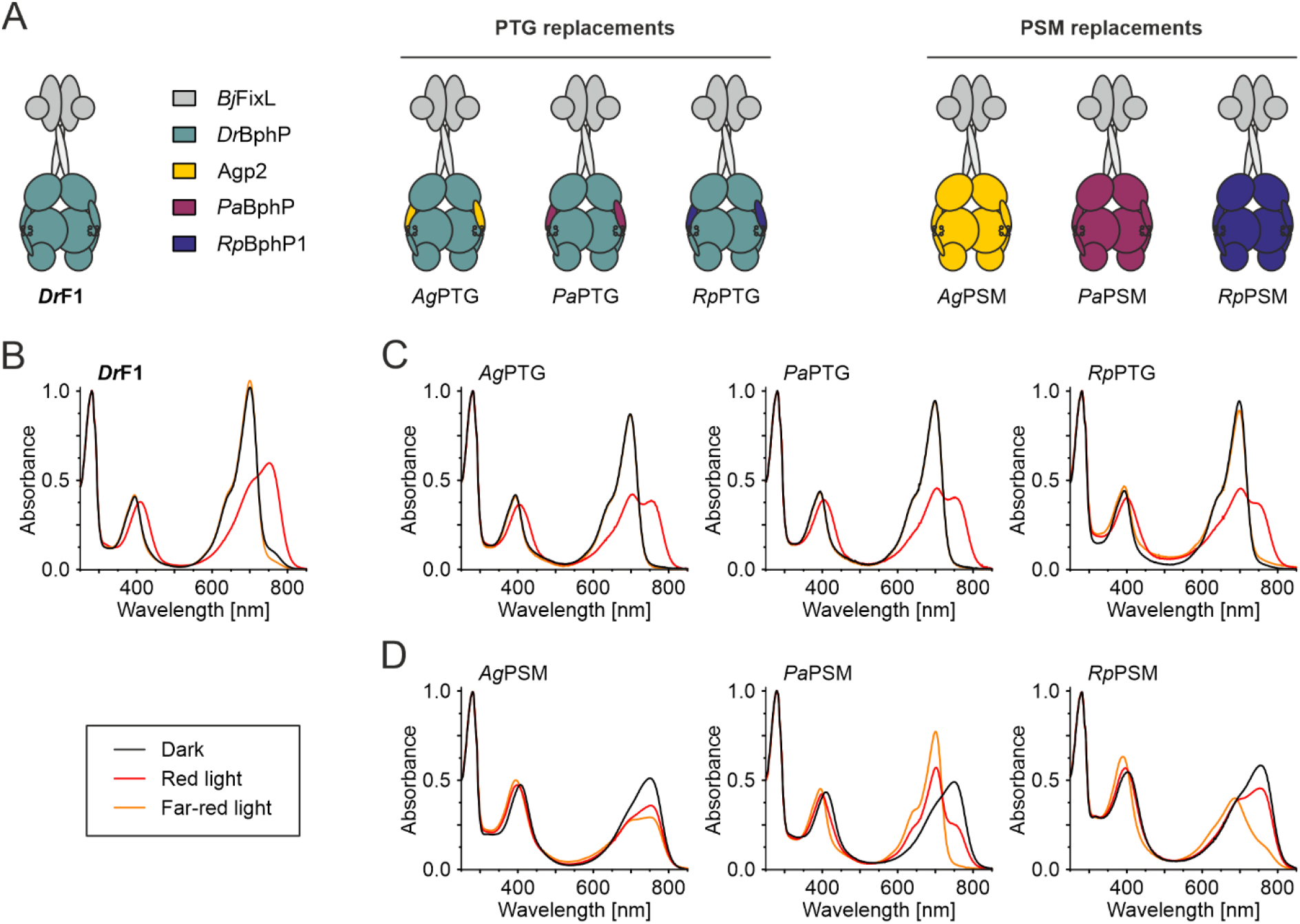
DrF1 bathy chimaeras and their absorption spectra. **(A)** Schematic presentation of DrF1 chimeras containing structural elements from three different bathy BphPs. Origins of the derived domain exchanges are colour coded as indicated, see Table S2 for domain boundary definitions. **(B)** Absorption spectra of DrF1^55^. **(C)** Absorption spectra of DrF1 chimaeras where the PHY-tongue (PTG) was exchanged for the corresponding bathy sequence. **(D)** DrF1 chimaeras where the entire bathy photosensory module (PSM) was introduced.

The most obvious way to generate a bathy variant of *Dr*F1 would be to replace its PSM with one of bathy origin. Therefore, we next decided to exchange the entire photosensory module of *Dr*F1 with the PSMs from the three bathy representatives. All three resulting constructs – *Ag*PSM, *Pa*PSM and *Rp*PSM (Fig. 3A) – exhibit bathy characteristics with resting state peak maxima at Pfr wavelengths of ∼750 nm (Fig. 3D). Although the PSM chimaeras occupy Pfr resting states, they differ in their Pr-state spectra. Only *Pa*PSM occupies a classic Pr state spectrum under far-red light (699 nm Pr Q-band maximum), whereas *Rp*PSM undergoes an incomplete hypsochromic shift of the Q-band in its Pr state (685 nm Pr maximum). Both these spectra resemble those published for *Pa*BphP and *Rp*BphP1 truncations^15,45^. In contrast, the *Ag*PSM chimaera barely shifts from the Pfr state at all, unlike reported for an Agp2 fragment^46^, which could be ascribed to the effects caused by artificial fusion to the *Bj*FixL OPM. Although the intensities of BV-specific Soret and Q-bands in *Pa*PSM are relatively low in the Pr state, the Pfr-state spectrum is less affected. As discussed by Meier *et al*., the p*K*_*a*_ of BV affects the intensity of the Q-band in Pr state^47^, implying that our chimeric substitutions affect the cofactor environment in some manner^47–54^. When the *Dr*BphP PHY-tongue is re-introduced into chimaeras with a bathy PSM (Fig. S4A), the constructs appear locked in the Pr state (Fig S4D), as also confirmed by urea denaturation (Fig. S5).

Consequently, we show here that bathy vs. prototypical BphP behaviour is in fact determined by the PSM, but it is not defined solely by the PHY-tongue or NTS. The former plays an integral role in establishing spectral properties in BphPs^37,44^, but BphPs likely require additional elements in the PSM to adopt bathy characteristics. The PHY-tongue has previously been proposed to be at a constant transition between closed and open states, both affected by and affecting equilibria between Pr- and Pfr-state BV conformations, PHY-tongue folding, as well as PSM and protein-wide structural arrangements^23,56,57^. It stands to reason that the definition of bathy or prototypical resting states might be determined by a similarly intricate interplay of various factors.

### OPM functionality is decoupled from light response in bathy chimaeras

The limited effect of PHY-tongue exchanges on spectral characteristics translates also to enzymatic functionality, as observed both *in vivo* in *Escherichia coli* cells utilising the pREDusk circuit^33^ (Fig. 4A), and *in vitro* on Phos-tag gels (Fig. 4B). In the pREDusk expression tool, net kinase activity and the resulting *Bj*FixJ phosphorylation lead to expression of the fluorescent marker protein *Ds*Red^58^. Phos-tag gels separate proteins according to their phosphorylation status, and the net kinase activity of *Dr*F1 variants is visible as an increased amount of phosphorylated *Bj*FixJ^36^. In both approaches, the results we observed for *Ag*PTG, *Pa*PTG and *Rp*PTG are all comparable to *Dr*F1 (Fig. 4), with high net kinase activity in the dark and under far-red light that is strongly suppressed by red light. Thus, *Dr*PSM is capable of controlling OPM functionality regardless of PHY-tongue origin. *In vitro* assays for the NTS exchanges also show similar behaviour to *Dr*F1 (Fig. S6C). These observations indicate that neither PHY-tongue nor NTS exchanges seem to affect *Dr*F1 enzymatic activity.

**Figure 4:**
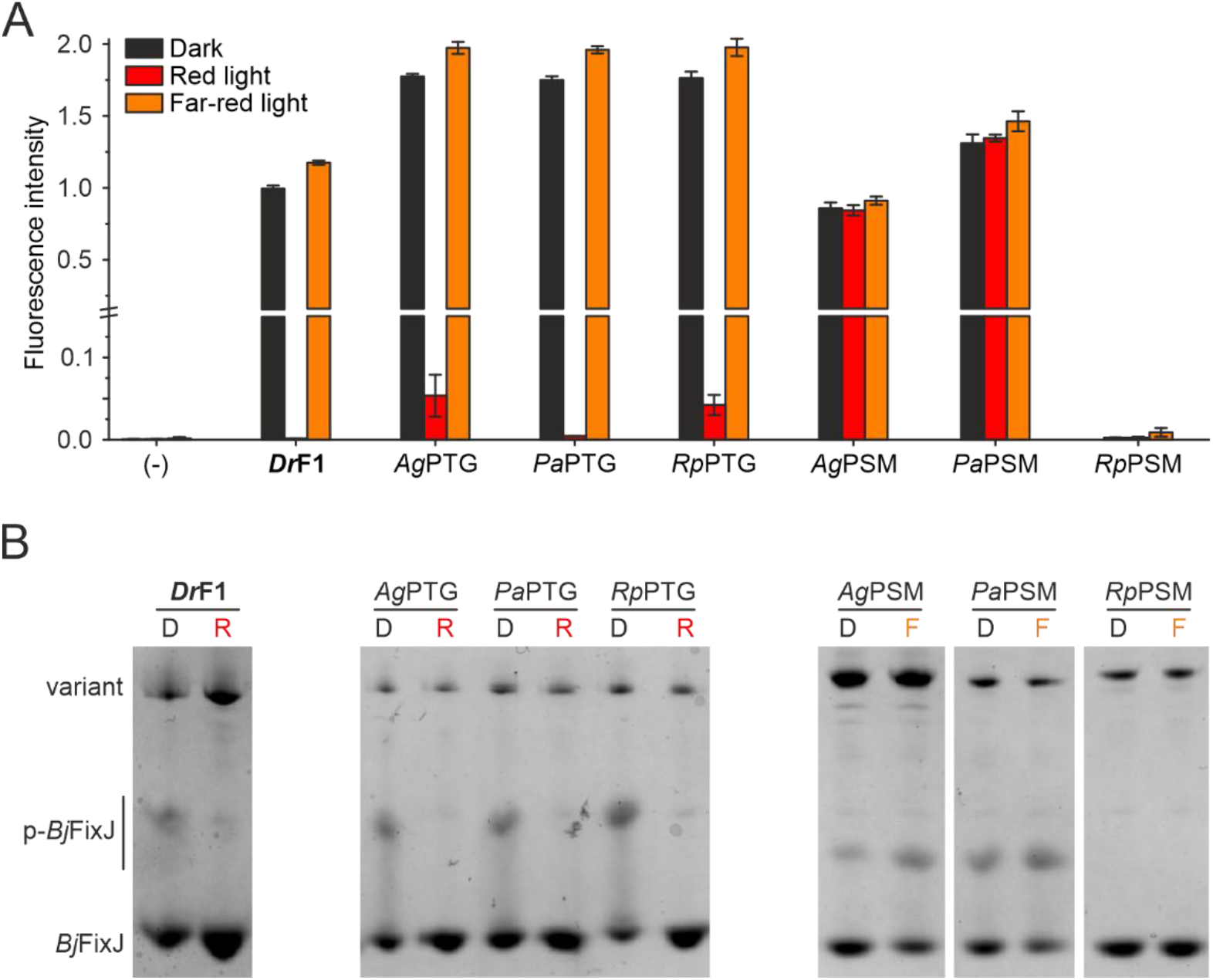
Histidine kinase activity analysis. **(A)** In vivo HK activity assay, where DsRed fluorescence (normalised to DrF1, dark) corresponds to net kinase activity of the DrF1 variant. Results are shown as mean ± SD of three biological repeats, see Figure S7A for individual biological repeats. pREDusk with the DsRed reporter gene replaced by a multiple cloning site (MCS) functions as negative control (-). **(B)** Phos-tag analysis of DrF1 variant activity in dark (D), in red light (R), or in far-red light (F). In the assay, phosphorylated BjFixJ (p-BjFixJ) migrates more slowly than the unphosphorylated one. See Figure S7B for full gels.

In contrast to the clear consensus between prototypical chimaeras and *Dr*F1-like HK activity regulation, the enzymatic activity of bathy variants (*Ag*PSM, *Pa*PSM and *Rp*PSM) was not regulated with light (Fig. 4) even though they appear bathy in their spectral behaviour (Fig. 3D). *Ag*PSM and *Pa*PSM feature similar *Ds*Red fluorescence levels under red light as under far-red light and dark, comparable to *Dr*F1 in the dark. We also observed the same effect in Phos-tag assays, where net kinase activity appears constantly high due to the loss of light control. *Rp*PSM, on the other hand, lacks all kinase activity. Introduction of *Dr*PTG into bathy PSM variants has no further effect on light regulation of enzymatic activity (Fig. S6A,B), concordant with the lack of spectral response (Fig. S4D).

In the case of *Rp*PSM, the lack of light control can be explained by the incompatibility between the PSM and the HK module, as *Rp*BphP1 originally features a PAS/PAC OPM^19^. In *Ag*PSM and *Pa*PSM, we assume that the lack of light control was due to an incompatible linker helix between the PSM and the effector HK. This coiled-coil linker region – also termed “neck” or “signalling helix” – connects the PSM and the OPM (Fig. 1A) and has repeatedly been discussed as a major regulatory element for effector activity by impacting orientation and/or degrees of freedom of the OPMs^7,24,55,59,60^. This can in turn affect accessibility of catalytic residues in *cis*-as well as *trans*-acting enzymatic effectors, like HKs^11^. The two main factors pertinent to controlling effector properties are the sequence as well as the length of the linker element^59^.

### Bathy BphPs favour certain sensor-effector linker lengths

Subsequently, the variations in linker length found across BphPs with known spectral properties drew our attention. In fact, linker length varies considerably less among BphPs with a HWE-HK than among those with a HisKA OPM^55^, as the majority of known bathy BphPs have HWE-HKs as their OPM (Fig. 2) with a linker length of –3 relative to *Dr*BphP (Fig. S8)^24^.

Here, we define linker length from PRXSF+13 (*Dr*BphP P482) to the catalytic histidine residue in the HK module (*Dr*BphP H532). As Bódizs et al. have previously pointed out, *Pa*BphP stands out among known bathy BphPs with a linker length of –10 residues relative to *Dr*BphP^24^ (Fig. S8). This linker length is found in only one additional characterised bathy BphP, *Rt*BphP2^61^, as well as in several other BphPs for which spectral properties have not been elucidated. Among the known bathy BphP HKs, *Pa*BphP and *Rt*BphP2 are the only ones with a HisKA output module, as opposed to the HWE-HK OPMs found for the other nine (Fig. S8B). Though no exclusive clustering is observed in phylogenetic trees of BphPs with known spectral properties (Fig. S1), a co-evolution of bathy BphPs and HWE-HK OPMs has been proposed in the past^48^. Presumably, the linker length restraints observed for bathy BphPs might be a by-product of that evolutionary relationship, or they reflect the relatively small number of known bathy BphPs.

To test whether matching the linker lengths of *Ag*PSM and *Pa*PSM to those found in known bathy BphPs would lead to light control, we generated two linker truncations: *Ag*PSM_–3_, corresponding to Agp2 and most known bathy BphPs, and *Pa*PSM_–10_, matching the *Pa*BphP linker (Fig. 5A). However, neither linker truncation shows notable light regulation in a pREDusk framework, and only *Ag*PSM_–3_ shows net kinase activity (Fig. 5B). This indicates that exchanging the HK OPM in bathy BphPs to *Bj*FixL alters the requirements for linker helix length in a yet uncharacterised way.

**Figure 5:**
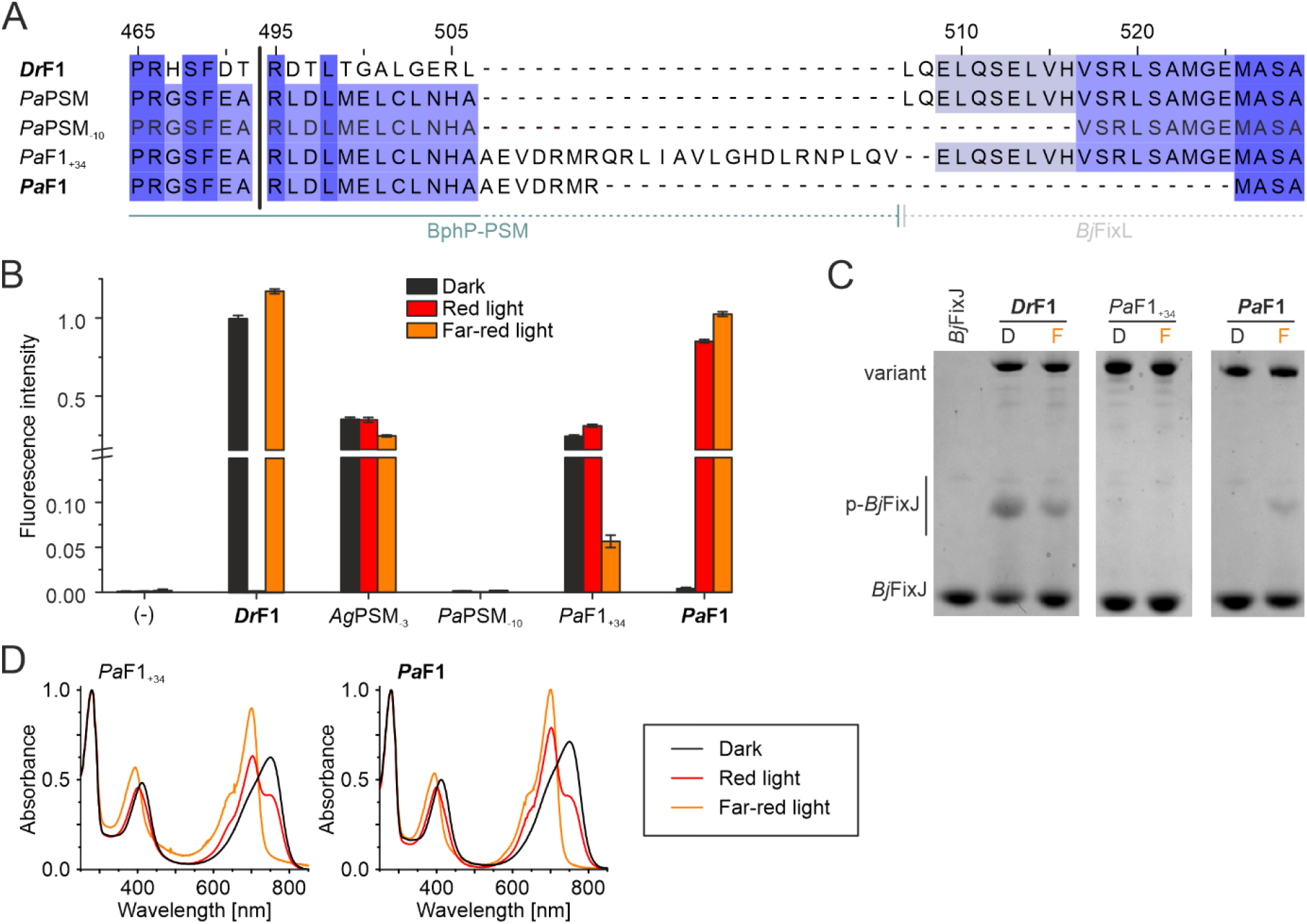
PaPSM linker length variants and PaF1. **(A)** Linker length comparison of DrF1^55^ and PaPSM linker deletions and PATCHY products. Coloured according to Jalview sequence ID (hues of blue correspond to ≥ 84%, ≥ 68%, ≥ 40%, < 40% residue conservation), numbering corresponds to DrBphP. **(B)** In vivo HK activity assay visualised by DsRed fluorescence, normalised to DrF1 (dark). Results are shown as mean ± SD of three biological repeats, see Figure S9B for individual biological repeats. pREDusk with the DsRed gene replaced by a multiple cloning site (MCS) functions as negative control (-). **(C)** Phos-tag analysis of the PaF1 variant activity in dark (D) or in far-red light (F). In the assay, where phosphorylated BjFixJ (p-BjFixJ) migrates more slowly in comparison. See Figure S9C for full gels. **(D)** Spectral properties of PaF1 and PaF1_+34_, both of which appear bathy.

### *Pa*F1: A far-red light activated HK component for the optogenetic toolbox

As the rational design approach did not result in functional bathy linker variants, we subjected *Pa*PSM to the PATCHY protocol, which generates a library of variants with different linkers^59^. We have successfully applied this method before to invert pREDusk function from a red light-repressed system to the red light-activated pDERusk tool^55^. PATCHY yielded two *Pa*PSM clones of interest, both of which contain an extended *Pa*BphP sequence compared to the original *Pa*PSM design (Fig. 5A). The first variant, termed “*Pa*F1_+34_”, has a long linker with 22 additional residues stemming from *Pa*BphP compared to *Dr*F1 (*Dr*F1 +22) and shows ∼5x down-regulation of activity by far-red light. This variant is a bathy HK with its kinase activity repressed by far-red light (Fig. 5B-D), which may serve as a template for a far-red light-repressed optogenetic expression tool in the future. The second candidate has a linker 12 residues shorter than *Pa*PSM and *Dr*F1 (*Dr*F1 -12). The protein itself is a photochromic bathy BphP (Fig. 5D) and shows strong kinase activity upregulation in response to far-red light both in an *E. coli* expression system (Fig. 5B) and as purified protein on Phos-tag gels (Fig. 5C). Due to its superior performance and far-red light-inducible behaviour, we chose this as our main target variant, naming it “*Pa*F1” and the corresponding optogenetic tool “pFREDusk”. In contrast to our previous pREDusk system^33^, this tool is activated by light and responds to far-red wavelengths (Fig. 6A).

**Figure 6:**
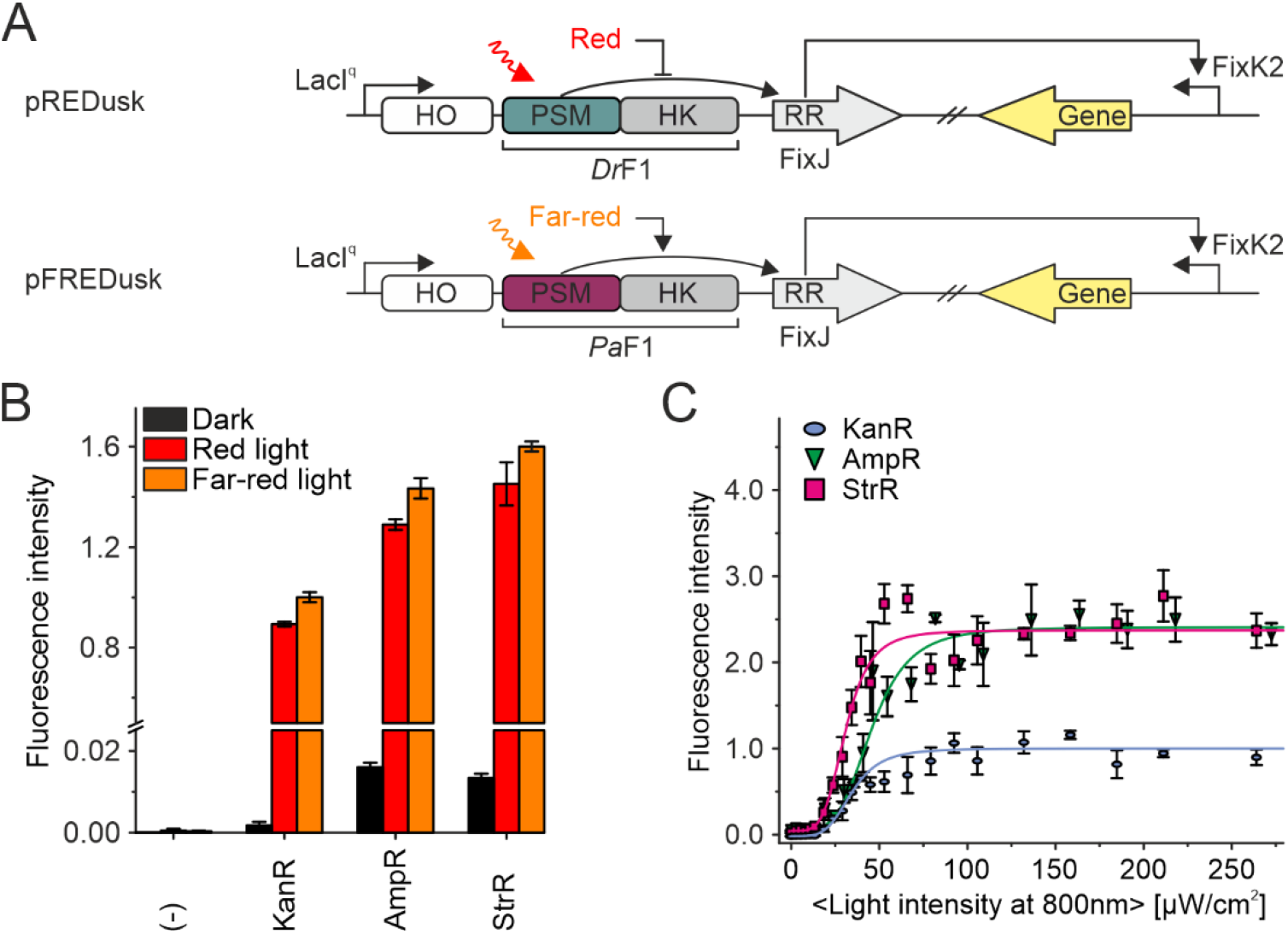
pFREDusk performance. **(A)** Schematic representation describing cellular pREDusk^33^ function (red light-repressed) compared to pFREDusk function (far-red light-activated). **(B)** PaF1-mediated in vivo HK activity. DsRed fluorescence is shown as mean ± SD of three biological repeats, normalised to KanR in far-red. See Figure S10B for individual biological repeats. pREDusk with the DsRed gene replaced by a multiple cloning site (MCS) functions as negative control (-). **(C)** DsRed production in bacteria harbouring pFREDusk versions with different antibiotic resistance in response to varied far-red light (800 nm) intensities, normalised to KanR endpoint. See Figure S10C for three biological repeats and Figure S10D for pFREDusk-KanR response to red light. Light intensities are averaged over the duty cycle as marked by angled brackets.

pFREDusk exhibits minimal background expression under non-inducing conditions and a several hundred-fold activity change upon illumination (Fig. 5B), which is comparable to the parent plasmid pREDusk^33^ as well as the NIR light-sensitive pNIRusk^47^. The bathy characteristic of *Pa*F1 in pFREDusk enables direct activation of this tool with far-red and red light. The dark reversion of *Pa*F1 is considerably faster than that of *Dr*F1, leading to full resting state within two hours (Fig. S3). Consequently, pFREDusk activation requires higher light intensities than pREDusk. pFREDusk also responds to red light, which is expected since red light creates a Pr/Pfr mixture, thus leading to a slightly smaller signal than far-red light illumination (Fig. 5B, Fig. 6B, Fig. S10B-D). Whereas *Dr*F1 is sensitive to both red and far-red light concordant with behaviour of *Dr*PSM^21^, only red light is able to repress pREDusk gene expression^33^.

To further widen the potential applicability of pFREDusk beyond kanamycin resistance (KanR), we also generated versions with streptomycin (StrR) and ampicillin (AmpR) resistance. Besides selection markers, different replication origins (ColE for KanR, CloDF13 for StrR and AmpR) allow circumvention of plasmid incompatibility with other optogenetic tools^62,63^. All three resulting pFREDusk versions respond to both far-red and red light with comparable upregulation of target gene expression (Fig. 6B). The lower overall expression levels of pFREDusk-KanR can be explained by the different origin of replication^64^. To gauge the far-red light sensitivity of the pFREDusk tools, we conducted light dose response experiments (Fig. 6C). The half-maximal light doses of pFREDusk-KanR, -StrR and -AmpR are (47 ± 11) µW cm^-2^, (36 ± 6) µW cm^-2^ and (42 ± 5) µW cm^-2^, respectively. In comparison to the originally reported pREDusk and pDERusk systems^33,55^, pFREDusk is one magnitude less sensitive to light. pFREDusk plasmids with all three antibiotic resistances have been made available in the Addgene repository (https://www.addgene.org), with the target gene replaced by a multiple cloning site (MCS) under accession IDs #229735 (pFREDusk-KanR-MCS), #229736 (pFREDusk-StrR-MCS), and #229737 (pFREDusk-AmpR-MCS).

## CONCLUSIONS

In this study, we introduce the pFREDusk tool that enables over 200-fold induction in target gene expression with far-red light in bacteria. By doing so, we provide an example for creating a far-red light-sensitive optogenetic system based on a well-studied prototypical one. Optogenetics is a field of increasing importance that uses light-sensitive proteins to control cellular functions^65^. Many optogenetic systems applied to eukaryotes are derived from prokaryotes; and bacterial applications are steadily increasing^2^. Red light-sensing systems attracting particular attention due to their potential for therapeutic and biotechnological applications. Red and far-red light at the NIR tissue transparency window^34,35^ penetrates deeper into tissues and bacterial cultures and with lower phototoxicity than shorter wavelengths, and BV, a breakdown product of haem, should be available in most mammalian tissues^66^. Although BphPs are capable of sensing far-red light as well as red light, the number of far-red light-regulated bacterial tools published to date remains sparse^67^. Also, far-red responding tools would show less cross-activation with other optogenetic tools of shorter wavelengths, *i*.*e*. controlled by blue light as shown for the pREDusk framework^33^, thus enabling wider and easier applicability to multichromatic optogenetic systems^66,68^. Induction of target effects is generally preferable to repression for most implementations of optogenetic systems. We expect the pFREDusk circuit to be of great interest for a wide variety of applications and research approaches, ranging from studies on the importance of individual bacterial genes to adaptation for control of gene expression in mammalian cells.

While developing the pFREDusk tool, we also demonstrate the complexity of bathy BphP characteristics, and the many structural elements involved in defining the BphP photoresponses. A total of three characteristics were found more strongly conserved within bathy BphPs than across BphPs in general: 1) a discrete length of the PHY-tongue, 2) a glycine in position PRXSF–8, and 3) HWE-HK OPMs. These characteristics, however, are not exclusively limited to bathy BphPs, nor present in all known bathy representatives. With a comparatively limited number of BphPs investigated spectroscopically, it would be interesting to gain better insight into phylogenetic, structural and functional relationships of bathy and prototypical BphPs by screening a larger number of naturally occurring BphPs. This would also facilitate the design of far-red light regulated artificial systems. pFREDusk and its actuator *Pa*F1 further underpin the importance of the linker element between the sensor and output module in photoreceptor signal integration and highlight the far-reaching potential of semi-rational design of optogenetic tools. We also illustrate the possibility of creating far-red light-regulated optogenetic tools from existing prototypical systems through PSM exchanges. A switch into bathy BphPs can open up new possibilities in several biotechnological fields.

## METHODS

### Protein preparation, expression, and purification

*Dr*F1 and *Dr*F1_–6b_ have been described previously^33,55^. Residue fusion points for individual chimaeras are listed in Table S2. Bathy PTG, PSM, and PSM-*Dr*PTG constructs were created from the pREDusk plasmid^33^ using PCR-based Gibson assembly, and subsequently cloned into pET-21b(+) vectors. NTS and NTS/PTG constructs were assembled from pET-21b(+) *Dr*F1, *Ag*PTG and *Pa*PTG. pREDusk *Ag*PSM_–3_ and *Pa*PSM_–10_ plasmids were produced using the QuikChange Lightning Site-Directed Mutagenesis Kit. All plasmids used in this study are listed in Table S3. For implementation of the PATCHY protocol^59^, pREDusk *Pa*PSM was linearised and the linker region extended (Table S2) by introducing a GeneStrand insert (Eurofins Genomics), including a *Ksp*AI restriction site and a 1-nucleotide frame shift between PSM and *Bj*FixJ. The extended construct was then subjected to the PATCHY protocol as described previously^55,59^. Following PCR with pooled forward and reverse primers (Table S4) created with a custom Python script (https://github.com/vrylr/PATCHY), PCR products were purified from agarose gels and digested with *Ksp*AI. Purified reaction products were phosphorylated with T4 polynucleotide kinase and subsequently ligated with T4 DNA ligase. After transformation into *E. coli* XL 10-Gold cells, transformants were screened for far-red light sensitivity.

For protein expression, plasmids were transformed into *E. coli* BL21-DE3 cells. LB medium was inoculated from overnight cultures, and main cultures were grown at 37°C until an OD_600_ of 0.5–0.8. After induction with 0.5 mM isopropyl-β-d-thiogalactopyranoside (IPTG), cultures were cooled to 20°C overnight. Harvested cells were lysed with an EmulsiFlex system in lysis buffer (20 mM Tris pH 8.0, 300 mM NaCl) and centrifuged (30 min, 47,850 xg), before the soluble fraction was applied to Ni^2+^-NTA affinity HisTrap columns (GE Healthcare). Apoproteins were eluted in purification buffer (50 mM NaPO pH 8.0, 150 mM NaCl) with increasing imidazole concentration (1–500 mM gradient) and incubated in a molar excess of biliverdin hydrochloride overnight on ice. The following day, holoproteins were further purified using size exclusion chromatography (SEC, HiLoad 26/600 Superdex 200 pg column, GE Healthcare) in SEC buffer (30 mM Tris pH 8.0). Purified protein was flash-frozen in liquid nitrogen and stored at –80°C.

### Spectroscopic characterisation

Proteins were diluted to 2.5 µM in buffer (30 mM Tris pH 8.0, 150 mM NaCl) and spectra recorded using an Agilent Cary 8454 UV-Visible spectrophotometer. Dark state spectra were taken directly after thawing and dilution under non-actinic conditions. Light-state spectra were acquired at room temperature after 1 min illumination with a red (662 nm; prototypical) or far-red (782 nm; bathy) laser. Spectra were normalised to *A*_280_ = 1.

### Phos-tag analysis

Kinase activity analysis using Phos-tag acrylamide assays was performed as described previously^69^. Briefly, protein was diluted to 0.75 mg mL^-1^ (*Ag*PSM, *Ag*PSM-*Dr*PTG, *Rp*PSM and *Rp*PSM-DrPTG due to low band visibility on the gels) or 0.375 mg mL^-1^ (all other constructs) and prototypical constructs were illuminated with a far-red laser (782 nm) for 10 s to ensure Pr population. After mixing with reaction buffer (25 mM Tris/HCl pH 7.8, 5 mM MgCl_2_, 4 mM β-mercaptoethanol, 5% ethylene glycol) and 1.5 mg mL^-1^ *Bj*FixJ, proteins were illuminated for 10 s with a red (662 nm; prototypical) or far-red (782 nm; bathy) laser. Subsequently, reactions were initiated by addition of 2 mM adenosine triphosphate (ATP). Samples were then incubated for 20 min at 25°C under inducing (657 nm LED, 3.2–7 mW cm^-2^ and 780 nm LED, 0.78–1.2 mW cm^-2^ for red and far-red light, respectively) or non-inducing conditions and phosphorylation reactions were stopped by addition of SDS-PAGE sample buffer containing β-mercaptoethanol and 1 mM ZnCl_2_. Samples were subsequently subjected to mobility shift detection using Zn^2+^-Phos-tag SDS-PAGE (Wako chemicals), using 9% gels containing 13 µM Phos-tag acrylamide and a constant current of 40 mA. PageBlue (ThermoScientific) was used for staining.

### Histidine kinase assay

*In vivo* HK activity assays were adjusted from Multamäki et al.^33,70^, see Figure S10A for a schematic representation. Bacteriophytochrome variants were cloned into the pREDusk plasmid (Addgene plasmid #188970) containing the *Ds*Red Express2 red-fluorescent reporter^58^. The plasmids were transformed into *E. coli* DH5α and overnight cultures in LB under kanamycin (Kan) selection (25 μg mL^-1^) stored in 25% glycerol at 80°C. Bacteria from glycerol stocks were then streaked on LB/Kan agar plates and incubated overnight at 37°C in darkness. LB/Kan liquid cultures were inoculated with single colonies and incubated under non-inducing illumination conditions (dark for *Ag*PSM, *Pa*PSM, *Rp*PSM, *Ag*PSM-*Dr*PTG, *Pa*PSM-*Dr*PTG, *Rp*PSM-*Dr*PTG, *Pa*PSM_–10_ and *Pa*F1; red for *Dr*F1, *Ag*PTG, *Pa*PTG and *Rp*PTG; far-red for *Ag*PSM_–3_ and *Pa*F1_+34_) for 18 h at 30°C and 225 rpm. The following day, the liquid cultures were diluted 1:100 into LB/Kan, transferred to 96-well culture plates (83.3924, Sarstedt) and incubated in dark, under red light or under far-red light, respectively, for 18 h at 750 rpm. Absorbance at 600 nm and reporter fluorescence (λ_EX_ 554 ± 5 nm / λ_EM_ 591 ± 10 nm) were measured with a Tecan Spark Multimode Microplate Reader. All fluorescence measurements were performed with constant gain. Background signal (LB/Kan containing wells) was reduced from both measurements and fluorescence readings were normalised to absorbance. Three biological replicates were cultivated for all variants in all three illumination conditions, with three technical replicates measured of each sample under each condition. pREDusk without a reporter gene served as negative control. Fluorescence signals for variants and negative control normalised to pREDusk in dark = 1 are plotted as mean ± s.d. For all light conditions, samples were put under constant illumination with light sources placed over the samples. For red light, an LED with λ_EMmax_ = 657 nm (SLS-0214-C, Mightex Systems) was used, and an LED with λ_EMmax_ = 780 nm (SLS-0310-C, Mightex Systems) for far-red light, with a minimum of 100 μW cm^-2^ each. Light intensities were regulated with a LED Driver Unit (LEDD1B T-Cube, Thorlabs) and measured using a handheld power meter (PM1000, Thorlabs). All incubation steps were performed protected from ambient light.

After deciding on the main target variant *Pa*F1, the corresponding pFREDusk tool was cloned under two different selection markers (streptomycin and ampicillin) and CloDF13 replication origin in a similar fashion to pREDusk^33^. Histidine kinase assays were performed for these versions as described above, but with different antibiotic selection (150 μg mL^-1^ ampicillin and 50 μg mL^-1^ streptomycin). Data analysis was performed as described above, normalised to pFREDusk-KanR under far-red light.

### Light-dose response measurements

The light-dose responses of the different pFREDusk systems were measured as described before^33,55^. Briefly, after transformation of the plasmids into *E. coli* CmpX13 cells ^71^, inoculated overnight cultures containing 5 mL LB/antibiotic (50 µg mL^-1^ kanamycin, 100 µg mL^-1^ streptomycin, 50 µg mL^-1^ carbenicillin) were incubated for 24 h at 30°C and 225 rpm under non-inducing conditions (darkness). The overnight cultures were diluted 100-fold in 20 mL LB/antibiotic and 200 µL aliquots were dispersed into 64 wells of black-walled microtiter plates with a clear bottom (µClear, Greiner BioOne). After sealing the microtiter plates with a gas-permeable membrane, the samples were illuminated from below with an Arduino-controlled 8×8 array of light-emitting diodes (LEDs)^72,73^ with peak wavelengths of 624 ± 8 nm (red) and 800 ± 18 nm (far-red), respectively. During an 18 h incubation at 37°C and 750 rpm, the samples were illuminated with short light pulses with a duty cycle of 1:2 (1 s far-red light + 1 s darkness). Light intensities were calibrated with a power meter (model 842-PE equipped with a 918D-UV-OD3 silicon detector, Newport). For each well the optical density at 600 ± 9 nm and the *Ds*Red fluorescence (λ_EX_ 554 ± 9 nm, λ_EM_ 591 ± 20 nm) was measured with a Tecan Infinite M200 Pro microtiter plate reader. The background control pFREDusk-KanR-MCS, which lacks the *Ds*Red reporter, was carried in parallel. Fluorescence was normalised to the optical density and the MCS background control was subtracted. Data were plotted as a function of averaged light intensity (derived from the duty cycle) and fitted to Hill binding isotherms with the Fit-o-mat software^74^. Displayed fluorescence intensity was normalised to far-red light illuminated pFREDusk-KanR endpoint = 1.

### Phylogenetic analysis

41 sequences of investigated BphPs were compiled from literature and supplemented with 363 sequences listed in the InterPro database^75^ with a PAS (IPR013654) – GAF (IPR003018) – PHY (IPR013515) architecture. The 404 entries were aligned using default ClustalO^76^ through the Jalview workbench^77^. For *k*pLogo frequence logos^78^, the entire alignment as well as all known bathy sequences were reduced to the PHY-tongue (*Dr*BphP R446 to R477). Residues between *Dr*BphP P456 and D457 were removed for the logo of all 404 sequences, since no empty residue positions are permitted for sequence logo compilation.

## SUPPORTING INFORMATION

This article contains the following supporting information:

Figures S1-S11 and Tables S1-S4 (phylogeny, thermal recoveries, spectral and activity analysis of additional constructs, technical replicates of main figures; BphPs listed, chimaera fusion points, genetic materials used, PATCHY primers; PDF)

Sequence list (XLSX) and alignments (PHY), PhosTag source data (XLSX) – ZIP

## ACKNOWLEDGEMENTS

We would like to thank Prof. Janne Ihalainen (University of Jyväskylä) for stimulating discussions and ideas, Dr Mervi Hyvönen (University of Jyväskylä) for cloning the pFREDusk-MCS plasmids for Addgene, and MSc Milla Ylimartimo (University of Jyväskylä) for the help with protein purification and spectroscopic analysis. The authors gratefully acknowledge support from the University of Jyväskylä Nanoscience Center.

