## Supporting Information for "Traits of bathy phytochromes and application to bacterial optogenetics"

| | |
|---|---|
| Figure S1 | S2 |
| Figure S2 | S4 |
| Figure S3 | S5 |
| Figure S4 | S6 |
| Figure S5 | S7 |
| Figure S6 | S9 |
| Figure S7 | S10 |
| Figure S8 | S12 |
| Figure S9 | S13 |
| Figure S10 | S15 |
| Table S1 | S17 |
| Table S2 | S18 |
| Table S3 | S19 |
| Table S4 | S20 |

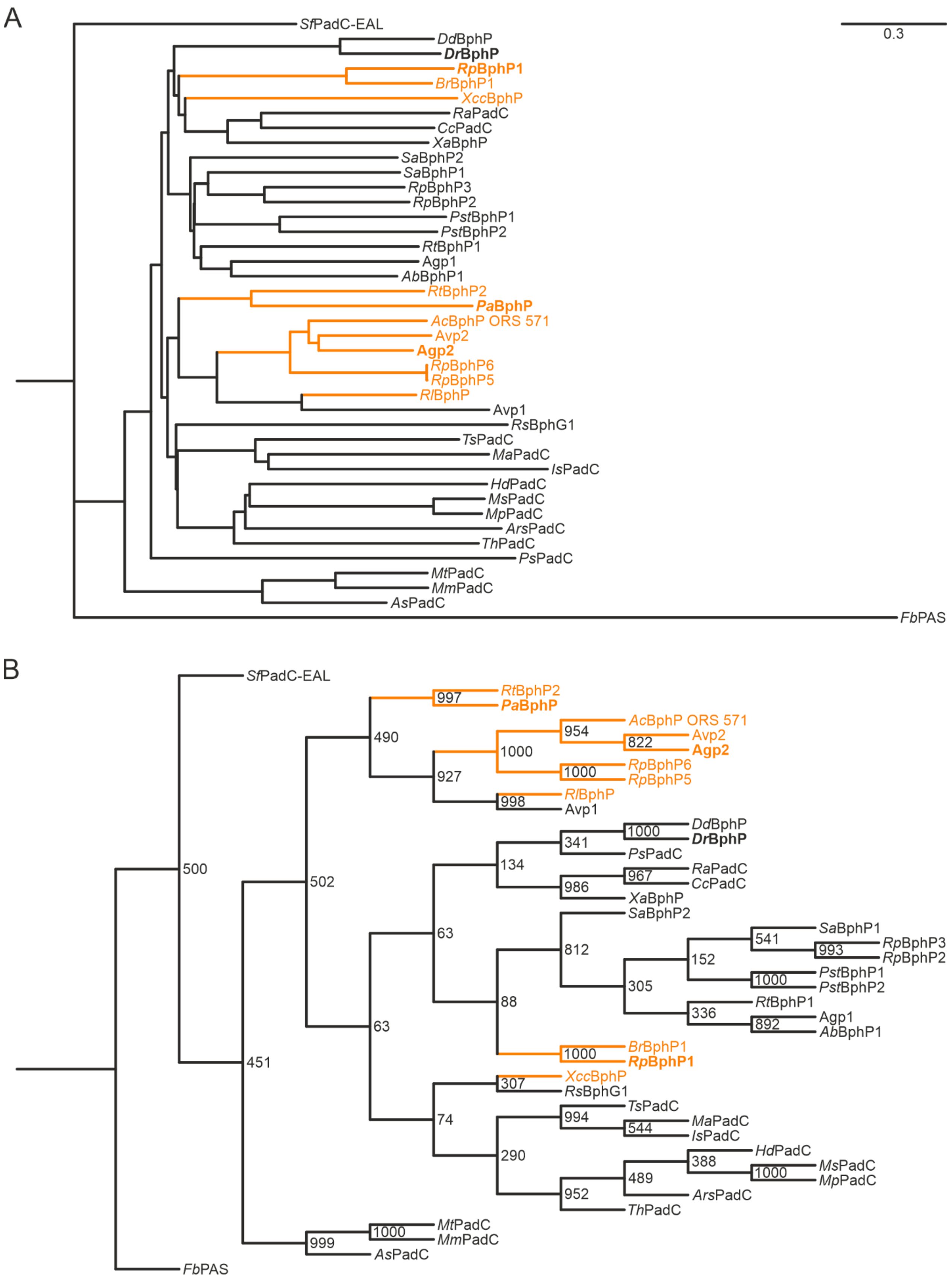
A
0.3
SfPadC-EAL
DdBphP
DrBphP
RpBphP1
BrBphP1
XccBphP
RaPadC
CcPadC
XaBphP
SaBphP2
SaBphP1
RpBphP3
RpBphP2
PstBphP1
PstBphP2
RtBphP1
Agp1
AbBphP1
RtBphP2
PaBphP
AcBphP ORS 571
Avp2
Agp2
RpBphP6
RpBphP5
RlBphP
Avp1
RsBphG1
TsPadC
MaPadC
IsPadC
HdPadC
MsPadC
MpPadC
ArsPadC
ThPadC
PsPadC
MtPadC
MmPadC
AsPadC
FbPAS
B
SfPadC-EAL
RtBphP2
PaBphP
AcBphP ORS 571
Avp2
Agp2
RpBphP6
RpBphP5
RlBphP
Avp1
DdBphP
DrBphP
PsPadC
RaPadC
CcPadC
XaBphP
SaBphP2
SaBphP1
RpBphP3
RpBphP2
PstBphP1
PstBphP2
RtBphP1
Agp1
AbBphP1
BrBphP1
RpBphP1
XccBphP
RsBphG1
TsPadC
MaPadC
IsPadC
HdPadC
MsPadC
MpPadC
ArsPadC
ThPadC
MtPadC
MmPadC
AsPadC
FbPAS
997
490
954
822
1000
1000
927
998
1000
341
134
967
986
500
502
63
812
541
993
152
1000
305
336
892
88
1000
63
451
307
74
994
544
290
388
1000
489
952
999
1000

***Figure S1:*** ***Phylogenetic trees of BphP PSM sequences with known spectral properties,*** *distance* ***(A)*** *vs. bootstrap tree* ***(B)****. BphP sequences were aligned with ClustalO default settings*[1] *(alignment available as supporting file, “Supp_known_aligned.phy”) and cut off after the PHY domain (last residue corresponds to DrBphP T499, before GALGERL motif). Trees were created using the PHYLIP 3.6 package (PROTDIST, FITCH)*[2]*. For the bootstrap tree, 1000 Jackknife subalignments were created using SEQBOOT and distance trees of subalignments were merged with CONSENSE. Trees were visualised with Figtree*[3]*. A PAS domain-containing protein from Flavobacterium bernardetii (FbPAS, NCBI protein ID WP_166125522) serves as outgroup. Known bathy BphPs are represented in orange. Low bootstrap values at some branching points highlight the complicated relationships across BphPs.*

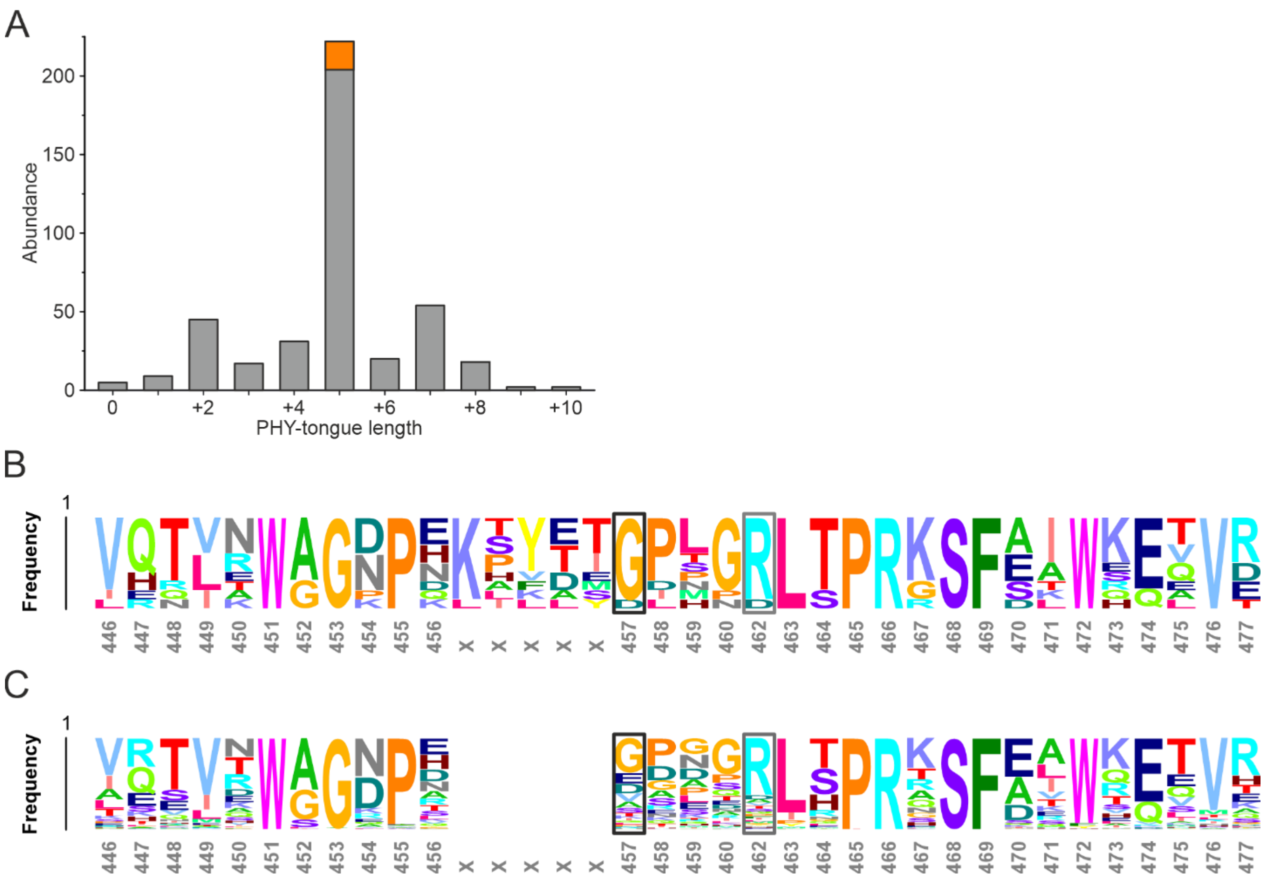


***Figure S2:*** ***PHY-tongue differences across BphPs. (A)*** *Absolute abundance of PHY-tongue lengths found in a ClustalO-aligned compilation of 404 BphP sequences. The tongue length is counted from WAG–5 to PRXSF+8 and given relative to DrBphP (32 residues). Tongue length of known bathy BphPs (+5) is marked in orange.* ***(B-C)*** *kpLogo*[4] *of 11 known bathy BphPs (B) vs. 404 BphP sequences (C). Numbering corresponds to DrBphP, the gap represents differences in tongue length. The glycine residue corresponding to DrBphP D457 is notably more conserved in bathy BphPs than in BphPs overall.*

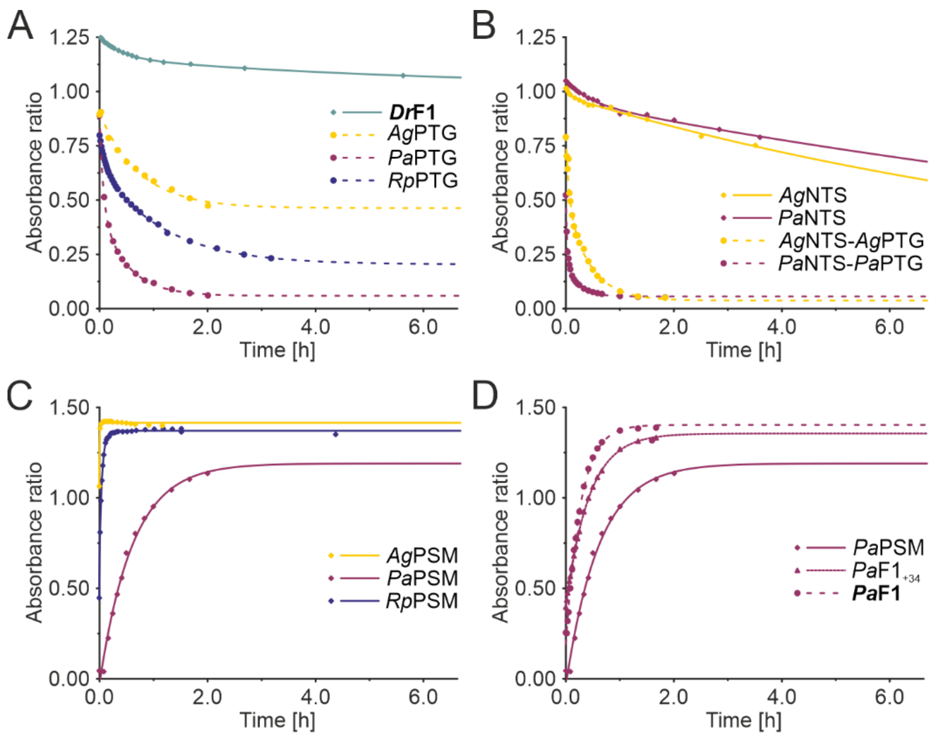


E

| Sample | exp | $A_{max/min}$ | $\tau_1$ (min) | $A_1$ (%) | $\tau_2$ (min) | $A_2$ (%) |
|---|---|---|---|---|---|---|
| ***Dr*F1** | 2 | 1.016 | 0.042 | 39 | 0.003 | 61 |
| *Ag*PTG | 1 | 0.464 | 0.023 | 100 | - | - |
| *Pa*PTG | 2 | 0.059 | 0.256 | 51 | 0.034 | 49 |
| *Rp*PTG | 2 | 0.204 | 0.154 | 23 | 0.014 | 77 |
| *Ag*NTS | 2 | 0.087 | 0.187 | 5 | 0.001 | 95 |
| *Pa*NTS | 2 | 0.001 | 0.039 | 9 | 0.001 | 91 |
| *Ag*NTS-*Ag*PTG | 2 | 0.038 | 0.274 | 39 | 0.039 | 61 |
| *Pa*NTS-*Pa*PTG | 2 | 0.056 | 0.660 | 65 | 0.075 | 35 |
| *Ag*PSM | 1 | 1.415 | 2.399 | 100 | - | - |
| *Pa*PSM | 1 | 1.190 | 0.027 | 100 | - | - |
| *Rp*PSM | 2 | 1.371 | 2.626 | 16 | 0.347 | 84 |
| *Pa*F1$_{+34}$ | 1 | 1.356 | 0.039 | 100 | - | - |
| ***Pa*F1** | 1 | 1.403 | 0.062 | 100 | - | - |

***Figure S3:*** ***Dark reversion of DrF1-bathy chimaeras,*** *plotted as the ratio of absorbance at 750/700 nm.* ***(A)*** *DrF1 and PHY-tongue (PTG)-exchange variants (Fig. 3A–C).* ***(B)*** *N-terminal segment (NTS)-exchange variants and NTS+PTG-exchange variants (Fig. S4A–C).* ***(C)*** *Photosensory module (PSM)- and PSM-DrPTG-exchange variants (Fig. 3A,D; Fig. S4A,D). The recovery kinetics of AgPSM-DrPTG, PaPSM-DrPTG and RpPSM-DrPTG could not be evaluated; all three constructs appear locked in Pr (Fig. S4D, Fig. S5).* ***(D)*** *PaPSM and its linker variants PaF1$_{+34}$ and PaF1 (Fig. 5D).* ***(E)*** *Time constants ($\tau_n$) and decay amplitudes ($A_n$) of the reversions. "exp" denotes the number of exponentials, and $A_{max/min}$ denotes the maximum absorption ratio at t = 0 min. The data was fitted using the formula* $\frac{A_{750}}{A_{700}}(t) = A_1 e^{-\frac{t}{\tau_1}} + A_2 e^{-\frac{t}{\tau_2}}$.

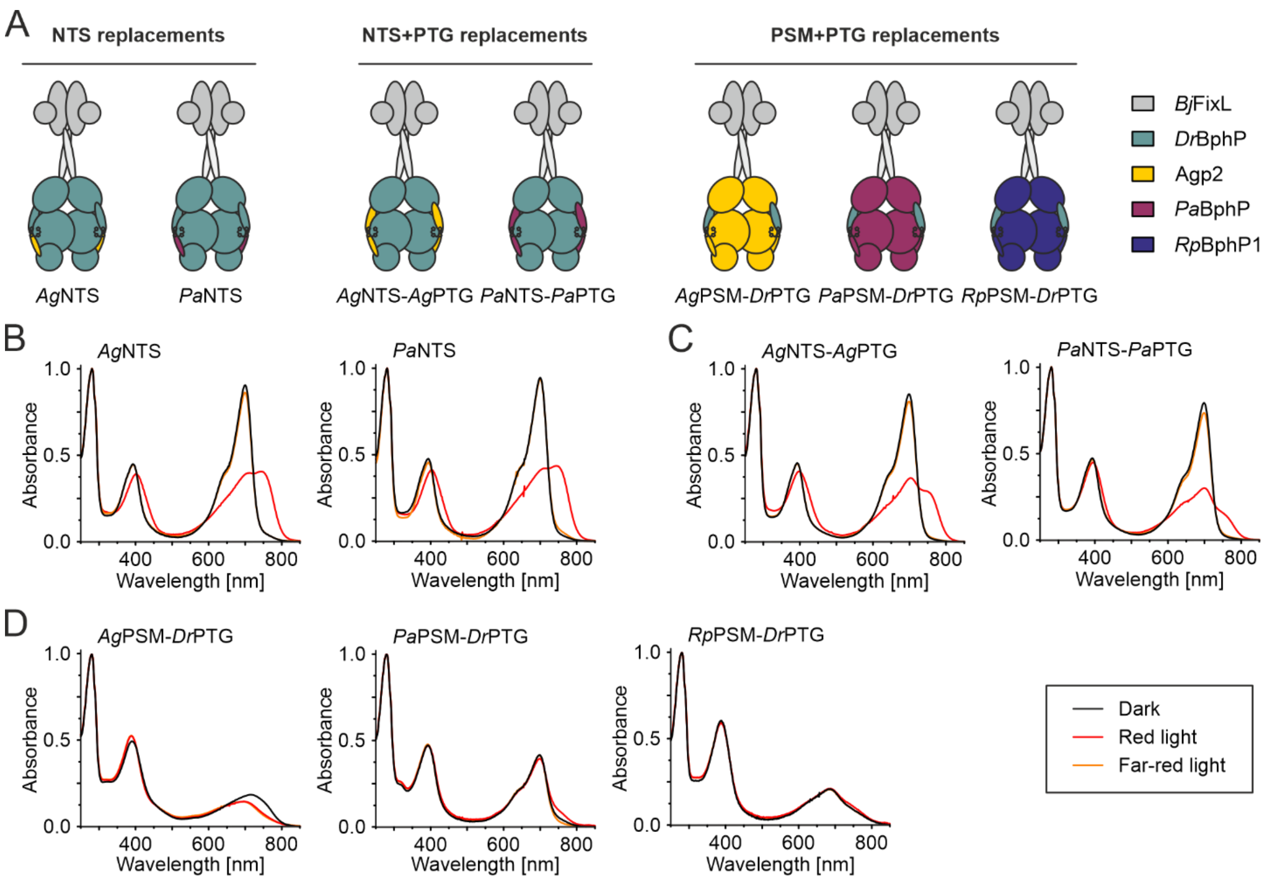


***Figure S4:*** ***Additional DrF1-bathy chimeras and their absorption spectroscopy.*** ***(A)*** *Schematic presentation of additional DrF1 chimeras containing structural elements from three different bathy BphPs. Origins of the derived domain exchanges are colour-coded as indicated, see Table S2 for domain boundary definitions.* ***(B)*** *Absorption spectra of DrF1 chimaeras where the N-terminal segment (NTS) was exchanged for the corresponding bathy sequence.* ***(C)*** *Absorption spectra of DrF1 chimeras where both NTS and PHY-tongue (PTG) were exchanged for the corresponding bathy sequence. Low Pfr contribution in the red light illuminated state is likely a result of notably fast dark reversion (Fig. S3B)* ***(D)*** *Absorption spectra of bathy PSM constructs where the PTG was exchanged back to the DrBphP sequence. The resulting constructs undergo only minor shifts upon illumination with either red or far-red light. The Q-band intensities of these three chimaeras are also notably low. Since BV peak intensities are similar for all chimaeras after urea denaturation (Fig. S5), the reduced Q-band extinction coefficients cannot be assigned to incomplete or faulty BV loading. All spectra are normalised to* $A_{280}$.

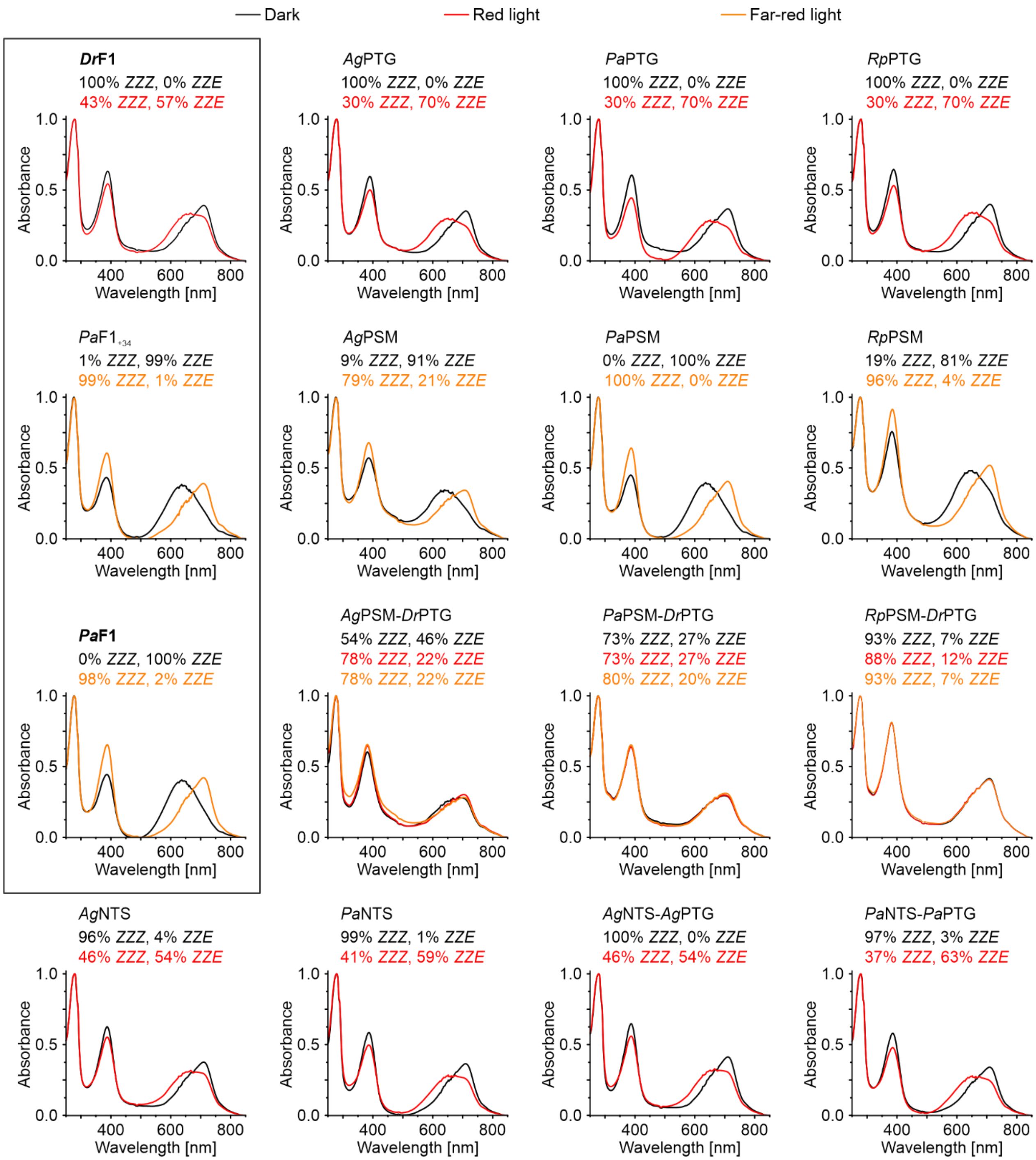


***Figure S5:*** ***UV/Vis analysis of bathy DrF1 chimaeras denatured with urea*** *at pH 3.0 to assess biliverdin (BV) isomerisation. 0.4 mg mL$^{1}$ protein were denatured after nonactinic / 3 min illuminated (662 / 782 nm) incubation and spectra were recoded using an Agilent Cary 8454 UVVisible spectrophotometer. Spectra are normalised to $A_{280}$ = 1 and scattering effects were subtracted. Percentages of BV in ZZZ and ZZE were calculated based on the assumption that the ZZE content in the Pr/Pfr*

*equilibrium reached after red light illumination equals 70% in DrPSM*[5]*, as previously described*[6]*. AgPSM-DrPTG, PaPSM-DrPTG and RpPSM-DrPTG are locked in the Pr state – in contrast to all other chimaeras that undergo Z/E isomerisation. Thus, DrPTG appears to lock the chromophore in a Pr like conformation within the otherwise bathy PSM environment, eliminating the canonical light response.*

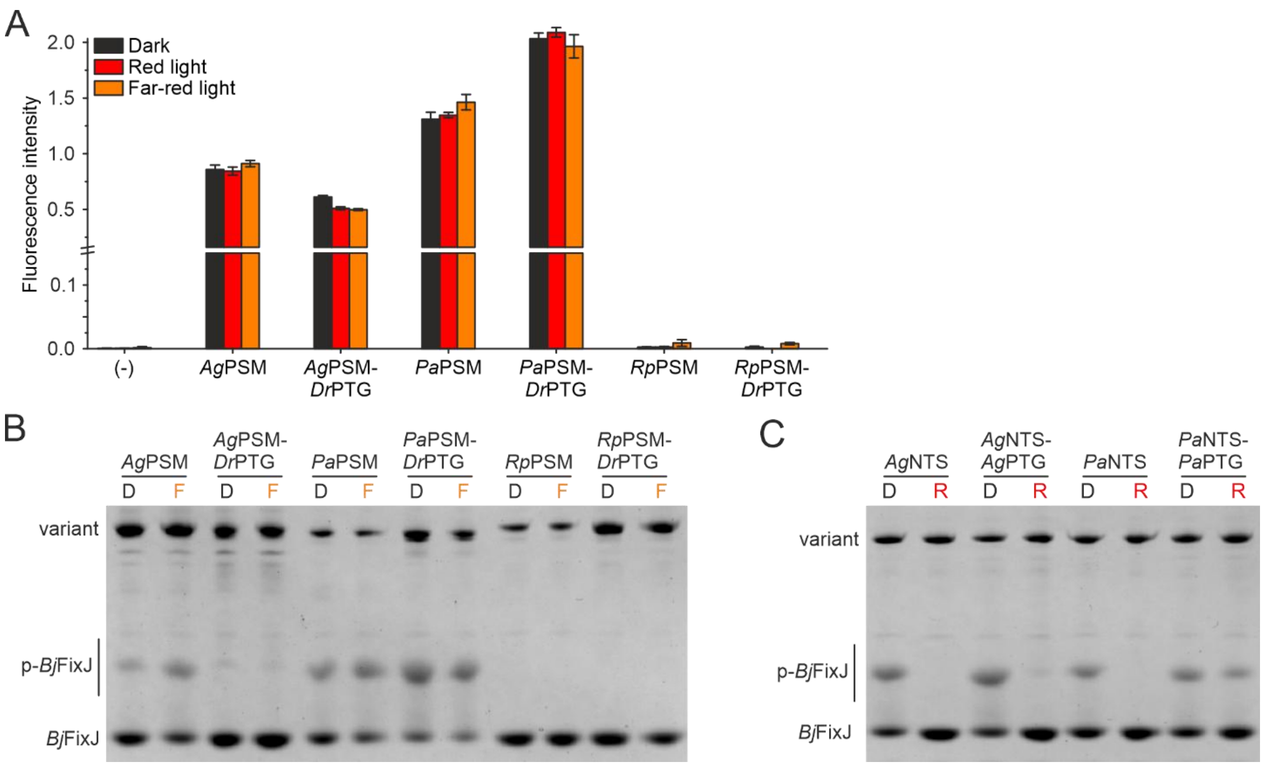


**Figure S6:** ***Activity assays of additional DrF1-bathy chimeras. (A)*** *Bacterial activity assay of PSM-DrPTG exchanges (Fig. S4A,D). The PSM exchanges from Figure 4 are also shown for comparison. pREDusk with the DsRed gene replaced by a multiple cloning site (MCS) functions as negative control (-).* ***(B)*** *Phos-tag activity assay PSM-DrPTG exchanges in dark (D) or in far-ref light (F). The PSM exchanges from Figure 4 are shown for comparison.* ***(C)*** *Phos-tag activity assay of N-terminal segment (NTS) exchanges (Fig. S4A,C), with and without corresponding PTG exchanges, in dark (D) or in red light (R). Samples were pre-illuminated with inducing light, mixed with ATP and incubated for 20 min under red (657 nm) / far-red (780 nm) light before separation on a polyacrylamide gel containing Phos-tag acrylamide, as previously described*[7].

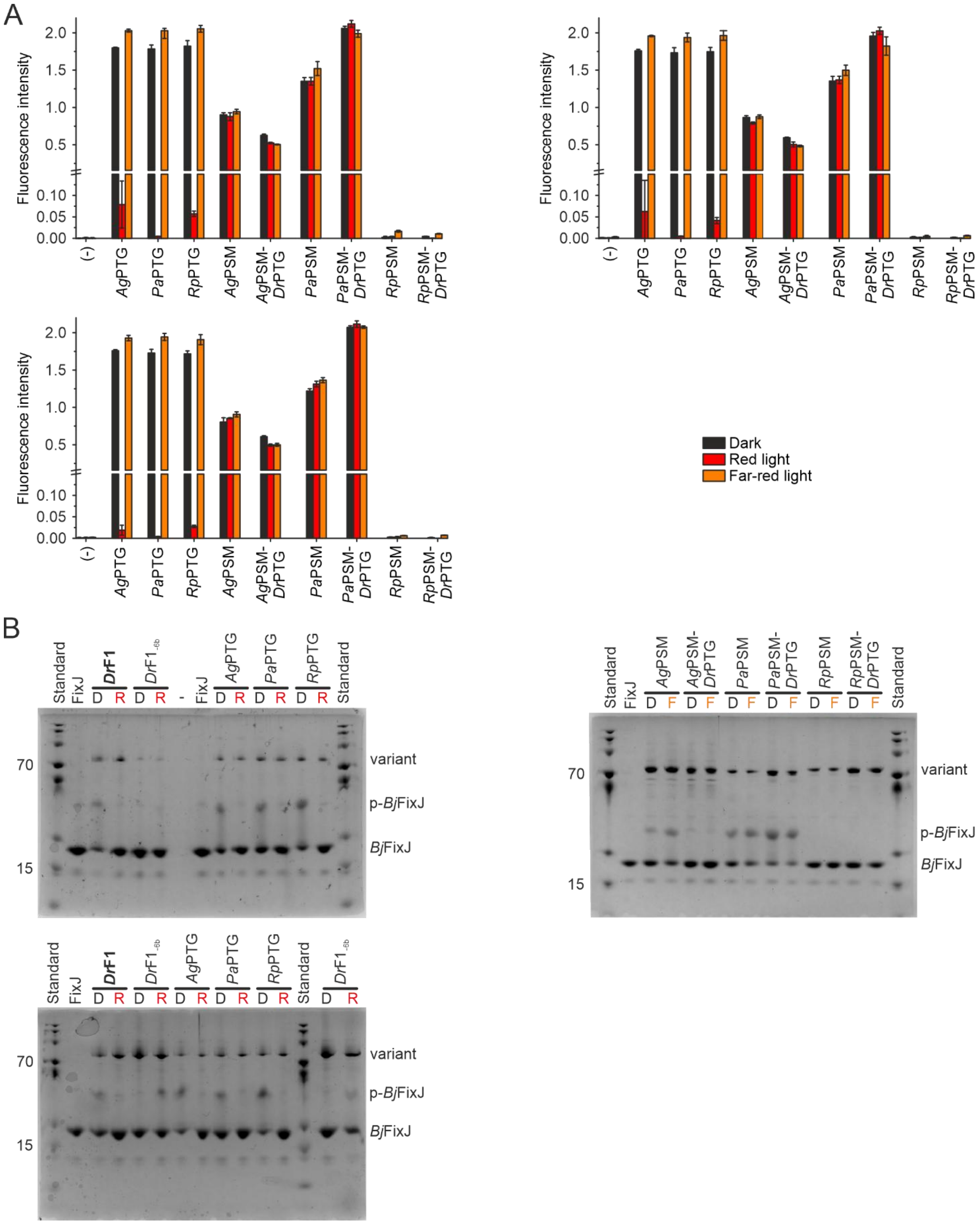


***Figure S7:*** ***In vivo & in vitro HK activity assays. (A)*** *Individual biological replicates of measurements depicted in Figure 4A and Figure S6. DsRed fluorescence (normalised to DrF1, dark) corresponds to histidine phosphorylation,*

*phosphotransfer to BjFixJ, and subsequent target gene expression. Individual biological repeats shown as mean ± SD of three technical repeats. pREDusk with the DsRed gene replaced by a multiple cloning site (MCS) functions as negative control (-).* ***(B)*** *Full gels as depicted in Figure 4B. Samples were pre-illuminated with inducing light, mixed with ATP and incubated for 20 min under red (657 nm) / far-red (780 nm) light before separation on a polyacrylamide gel containing Phos-tag acrylamide, as previously described*[7]*. PageRuler Plus Prestained Protein Ladder for reference (kDa) – BphP samples have ~80 kDa mass, BjFixJ has ~23 kDa.*

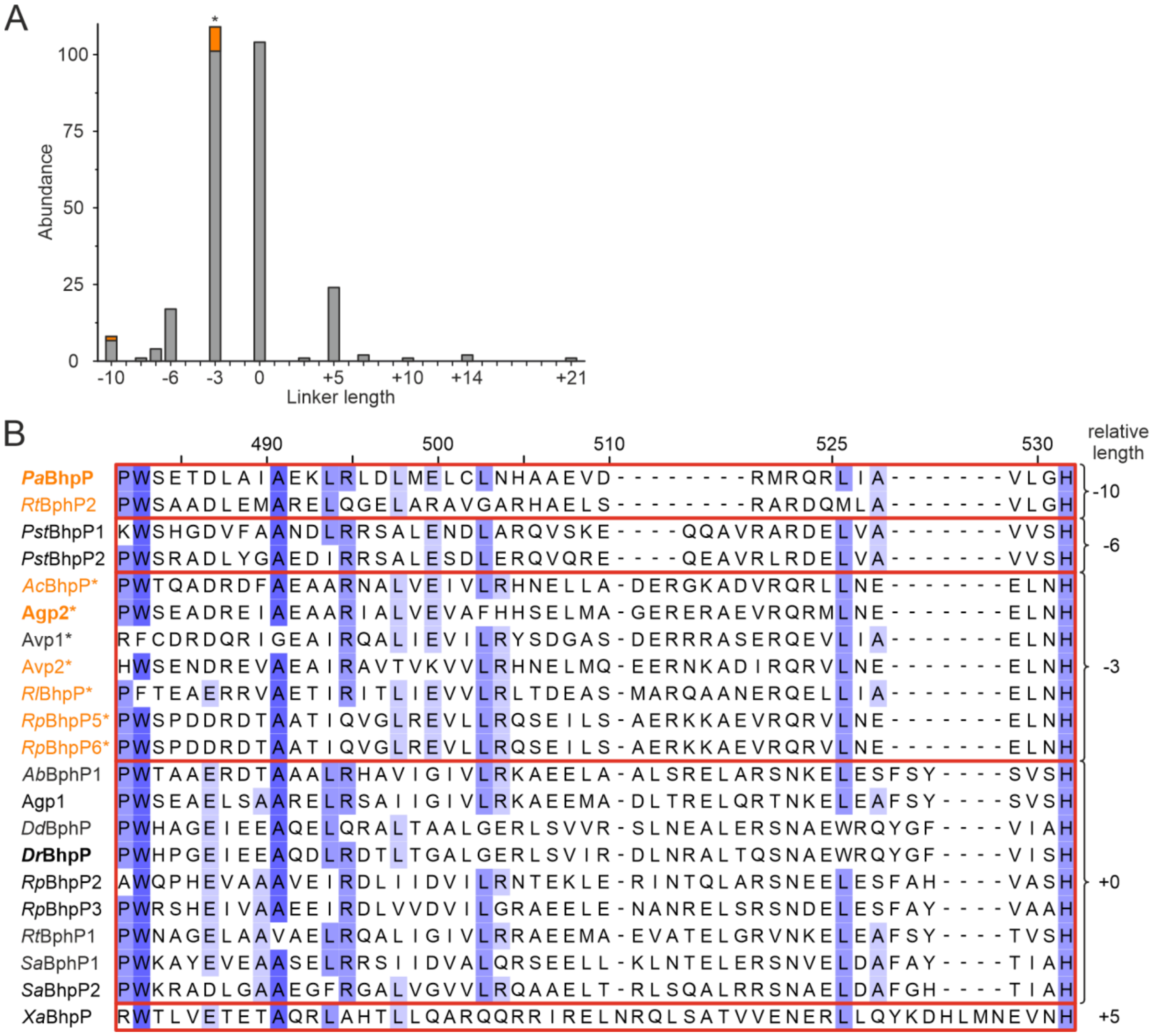


***Figure S8:*** ***Linker lengths across BphPs. (A)*** *Absolute abundance of linker lengths in a total of 275 published BphPs with a HK or HWE-HK OPM, linker counted from the conserved PW motif (PRXSF+13, DrBphP P482) to the catalytic His residue (DrBphP H532) and given relative to DrBphP (51 residues). 107 are HWE-HKs (*), 106 of which have a linker length of 48 residues. Known bathy BphPs are represented in orange.* ***(B)*** *Alignment coloured according to Jalview sequence ID (hues of blue correspond to ≥ 84%, ≥ 68%, ≥ 40%, < 40% residue conservation), numbering corresponds to DrBphP. 404 BphP sequences were aligned with ClustalO default settings[1] (full alignment available as supporting file, "Supp_sequences_aligned.phy") and manually curated. HWE-HKs are marked with *, bathy BphPs are represented in orange.*

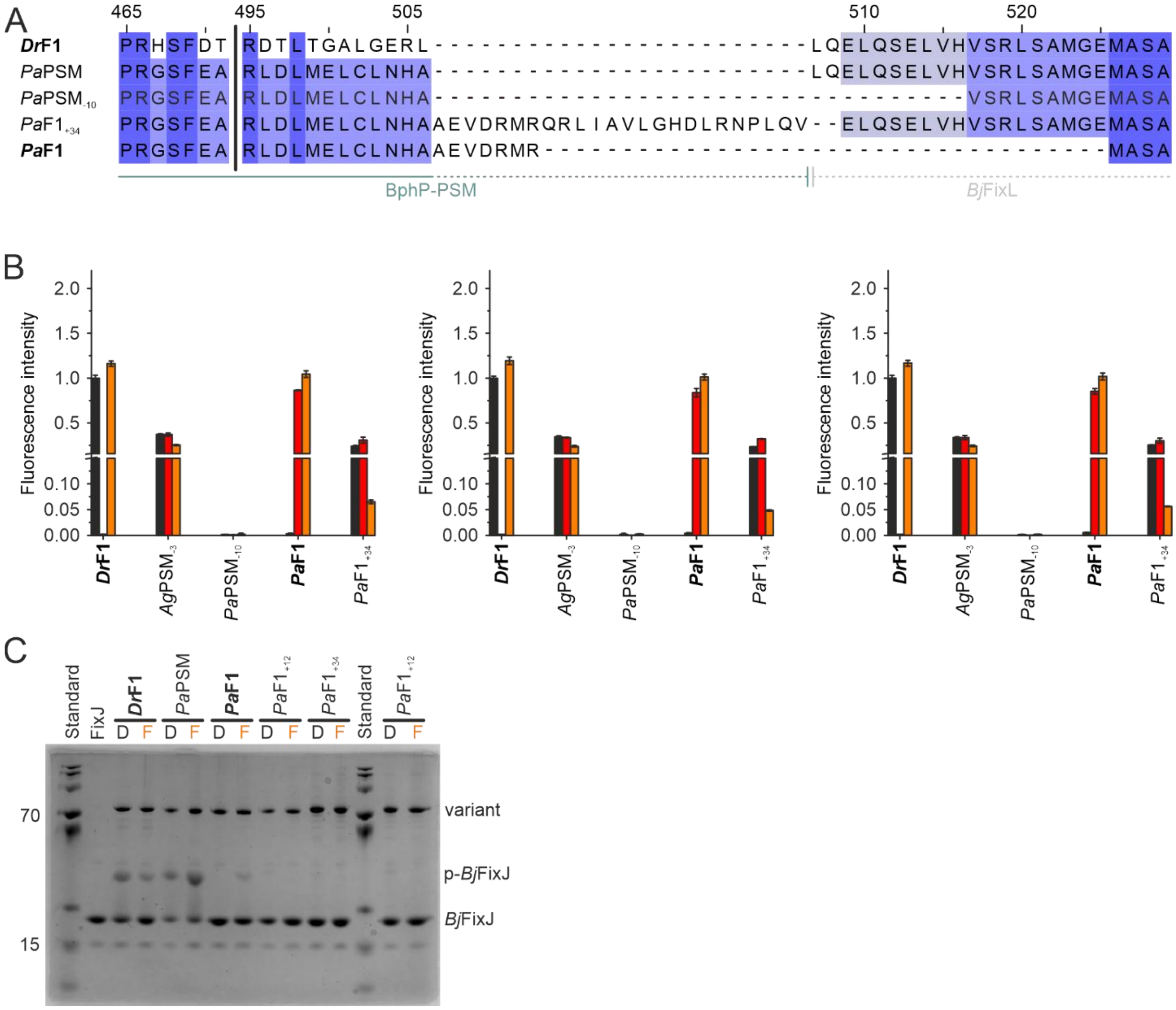


*Figure S9:* ***PaF1 activity in vitro and in vivo. (A)*** *Linker length comparison of AgPSM and PaPSM linker deletions. Coloured according to Jalview sequence ID (hues of blue correspond to ≥ 84%, ≥ 68%, ≥ 40%, < 40% residue conservation), numbering corresponds to DrBphP.* ***(B)*** *Individual biological replicates of measurements depicted in Figure 5B. DsRed fluorescence (normalised to DrF1, dark) corresponds to histidine phosphorylation, phosphotransfer to BjFixJ, and subsequent target gene expression. Individual biological repeats shown as mean ± SD of three technical repeats. pREDusk with the DsRed gene replaced by a multiple cloning site (MCS) functions as negative control (-).* ***(C)*** *Full gel as depicted in Figure 5C. Samples were pre-illuminated with inducing light, mixed with ATP and incubated for 20 min under red (657 nm) / far-red (780 nm) light before separation on a polyacrylamide gel containing Phos-tag acrylamide, as previously described[7].*

*PageRuler Plus Prestained Protein Ladder for reference (kDa) – BphP samples have ~80 kDa mass, BjFixJ has ~23 kDa.*

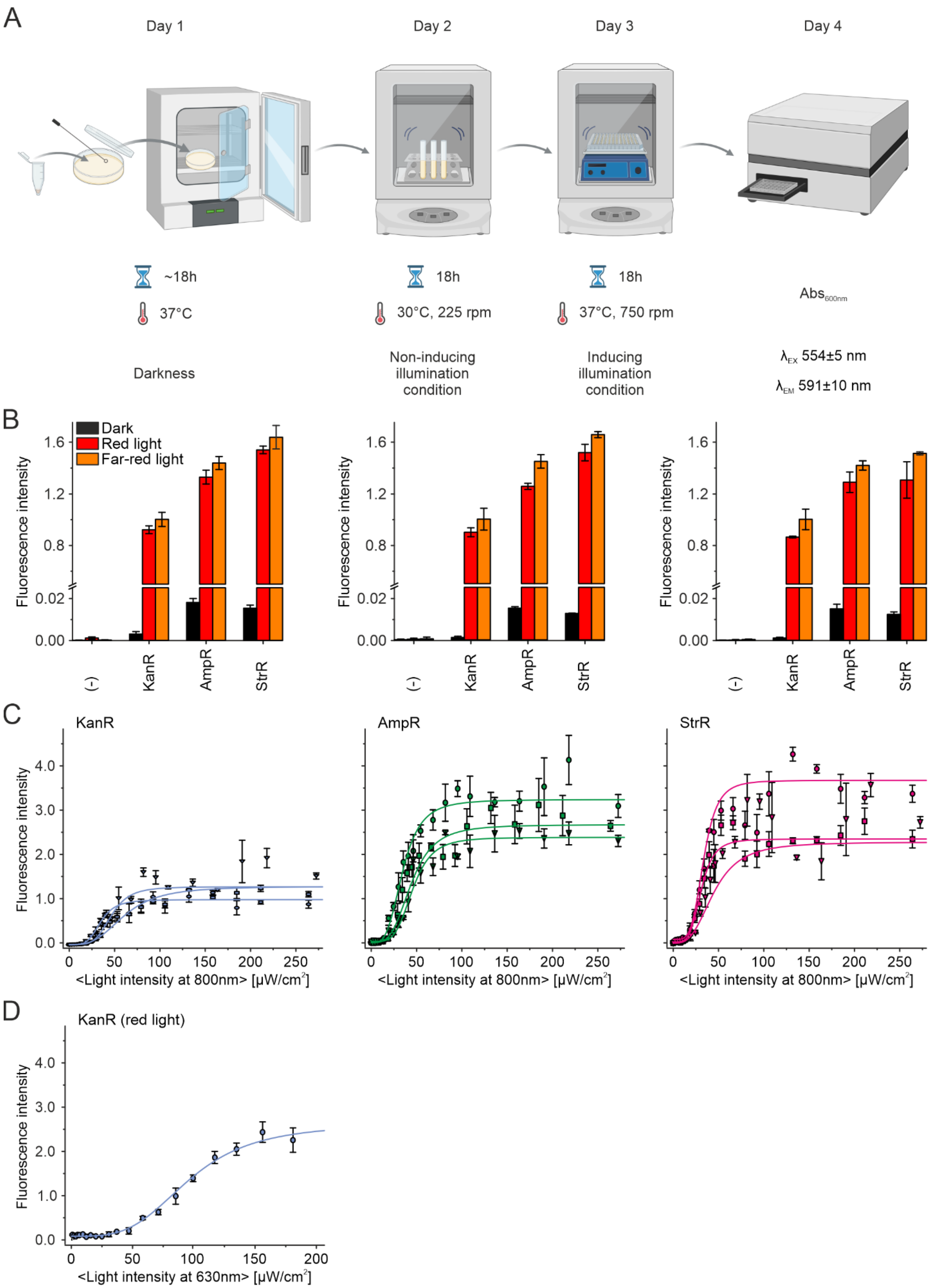
A
Day 1
Day 2
Day 3
Day 4
~18h
37°C
18h
30°C, 225 rpm
18h
37°C, 750 rpm
Abs600nm
Darkness
Non-inducing illumination condition
Inducing illumination condition
λEX 554±5 nm
λEM 591±10 nm
B
Dark
Red light
Far-red light
Fluorescence intensity
(-)
KanR
AmpR
StrR
C
KanR
AmpR
StrR
<Light intensity at 800nm> [μW/cm2]
D
KanR (red light)
<Light intensity at 630nm> [μW/cm2]

***Figure S10:*** ***Assessment of pFREDusk as an optogenetic tool. (A)*** *Schematic representation of in vivo kinase activity workflow. Depiction of the 4-day protocol, adjusted from Multamäki et al.*[8]*. The figure was created with BioRender.com.* ***(B)*** *Individual biological replicates of in vivo HK activity measurements depicted in Figure 6B. DsRed fluorescence corresponds to histidine phosphorylation, phosphotransfer to BjFixJ, and subsequent target gene expression. Individual biological repeats shown as mean ± SD of three technical repeats, normalised to KanR in far-red. pREDusk with the DsRed gene replaced by a multiple cloning site (MCS) functions as negative control (-).* ***(C)*** *Individual replicates of light sensitivity measurements depicted in Figure 6C. DsRed production in bacteria harbouring pFREDusk with KanR (blue), AmpR (green) and StrR (pink) in response to varied far-red light (800 nm) intensities. Fluorescence intensity normalised to far-red light illuminated KanR endpoint. Light intensities are averaged over the duty cycle as marked by angled brackets.* ***(D)*** *DsRed production in bacteria harbouring pFREDusk with KanR in response to varied red light (630 nm) intensities, fluorescence intensity normalised to far-red light illuminated KanR endpoint.*

***Table S1:*** ***Characterised BphPs*** *with known spectral properties as aligned in Figure 2 and Figure S8. Identifier as used in literature, parent organism, NCBI protein accession number and OPM.*

| **ID** | **Species** | **NCBI** | **OPM** |
|---|---|---|---|
| *Ab*BphP1 | *Azospirillum brasilense* | WP_040138077 | HK |
| *Ac*BphP ORS 571 | *Azorhizobium caulinodans* | WP_012169446 | HWE-HK |
| Agp1 | *Agrobacterium tumefaciens* F2 | EGP56260 | HK |
| Agp2 | *Agrobacterium tumefaciens* | WP_174008917 | HWE-HK |
| Avp1 | *Allorhizobium ampelinum* | WP_012650515 | HWE-HK |
| Avp2 | *Allorhizobium ampelinum* | WP_015916920 | HWE-HK |
| *Br*BphP1 | *Bradyrhizobium sp.* ORS 278 | WP_011924736 | PAC |
| *Dd*BphP | *Deinococcus deserti* | WP_012695249 | GHKL |
| *Dr*BphP | *Deinococcus radiodurans* | WP_010889310 | HK |
| *Is*PadC | *Idiomarina sp.* A28L | WP_007419415 | DGC |
| *Ma*PadC | *Marinimicrobium agarilyticum* | WP_027329460 | DGC |
| *Mp*PadC | *Marinobacter persicus* | WP_091706258 | DGC |
| *Pa*BphP | *Pseudomonas aeruginosa* | WP_034073960 | HK |
| *Pst*bphP1 | *Pseudomonas syringae* pv. *tomato* | KPY93576 | HK |
| *Pst*bphP2 | *Pseudomonas syringae* pv. *tomato* | WP_011104130 | HK |
| *Rl*BphP | *Rhizobium leguminosarum* | WP_184697183 | HWE-HK |
| *Rp*BphP1 | *Rhodopseudomonas palustris* | WP_119846077 | PAS/PAC |
| *Rp*BphP2 | *Rhodopseudomonas palustris* | WP_011158562 | HK |
| *Rp*BphP3 | *Rhodopseudomonas palustris* | WP_011158563 | HK |
| *Rp*BphP5 | *Rhodopseudomonas palustris* | WP_119019282 | HWE-HK |
| *Rp*BphP6 | *Rhodopseudomonas palustris* | WP_011156523 | HWE-HK |
| *Rs*BphG1, *Rhs*PadC-EAL | *Rhodobacter sp.* JA431 | WP_097082434 | DGC-EAL |
| *Rt*BphP1 | *Ramlibacter tataouinensis* TTB310 | AEG93643 | HK |
| *Rt*BphP2 | *Ramlibacter tataouinensis* TTB310 | WP_013902229 | HK |
| *Sa*BphP1 | *Stigmatella aurantiaca* | WP_002612494 | HK |
| *Sa*BphP2 | *Stigmatella aurantiaca* | WP_002609371 | HK |
| *Ts*PadC | *Thioalkalivibrio sp.* ALMg3 | WP_026331574 | DGC |
| *Xa*BphP | *Xanthobacter autotrophicus* Py2 | ABS67686 | HK |
| *Xcc*BphP | *Xanthomonas campestris* | WP_011270102 | PAS |

***Table S2:*** ***Residue numbers of fusion points for bathy DrF1 chimaeras introduced in this study.*** *PSM = photosensory module; NTS = N-terminal segment; PTG = PHY-tongue; OPM = output module. PaPSM ext. was the template construct for PATCHY PCR.*

<table>
<tr><th>Construct</th><th>PSM</th><th>NTS/PTG</th><th>OPM</th></tr>
<tr><td>AgPSM</td><td>Agp2 4–500</td><td rowspan="3">—</td><td rowspan="3">BjFixL 266–505</td></tr>
<tr><td>PaPSM</td><td>PaBphP 1–497</td></tr>
<tr><td>RpPSM</td><td>RpBphP1 1–506</td></tr>
<tr><td>AgPTG</td><td rowspan="3">DrBphP 1–506</td><td>Agp2 435–471</td><td rowspan="3">BjFixL 266–505</td></tr>
<tr><td>PaPTG</td><td>PaBphP 432–468</td></tr>
<tr><td>RpPTG</td><td>RpBphP1 441–477</td></tr>
<tr><td>AgPSM-DrPTG</td><td>Agp2 4–500</td><td rowspan="3">DrBphP 446–477</td><td rowspan="3">BjFixL 266–505</td></tr>
<tr><td>PaPSM-DrPTG</td><td>PaBphP 1–498</td></tr>
<tr><td>RpPSM-DrPTG</td><td>RpBphP1 1–506</td></tr>
<tr><td>AgNTS</td><td rowspan="4">DrBphP 1–506</td><td>Agp2 2–11</td><td rowspan="4">BjFixL 266–505</td></tr>
<tr><td>PaNTS</td><td>PaBphP 2–18</td></tr>
<tr><td>AgNTS-AgPTG</td><td>Agp2 2–11, 435–471</td></tr>
<tr><td>PaNTS-PaPTG</td><td>PaBphP 2–18, 432–468</td></tr>
<tr><td>AgPSM$_{-3}$</td><td>Agp2 4–500</td><td rowspan="2">—</td><td>BjFixL 269–505</td></tr>
<tr><td>PaPSM$_{-10}$</td><td>PaBphP 1–497</td><td>BjFixL 276–505</td></tr>
<tr><td>PaPSM ext.</td><td>PaBphP 1–523</td><td>—</td><td>BjFixL 262–505</td></tr>
<tr><td>PaF1</td><td>PaBphP 1–504</td><td>—</td><td>BjFixL 285–505</td></tr>
<tr><td>PaF1$_{+34}$</td><td>PaBphP 1–521</td><td>—</td><td>BjFixL 268–505</td></tr>
</table>

***Table S3:*** ***Genetic materials used in this study.*** *Antibiotic resistances: Kan = kanamycin; Amp = ampicillin; Str = streptomycin.*

| Construct | Parent plasmid | Insert | AntibioticR | Reference |
|---|---|---|---|---|
| pREDusk | pREDusk-*Ds*Red | *Dr*F1 | Kan | [8] |
| pDERusk | pREDusk-*Ds*Red | *Dr*F1$_{-6b}$ | Kan | [9] |
| — | pREDusk-*Ds*Red | *Ag*PSM | Kan | — |
| — | pREDusk-*Ds*Red | *Pa*PSM | Kan | — |
| — | pREDusk-*Ds*Red | *Rp*PSM | Kan | — |
| — | pREDusk-*Ds*Red | *Ag*PTG | Kan | — |
| — | pREDusk-*Ds*Red | *Pa*PTG | Kan | — |
| — | pREDusk-*Ds*Red | *Rp*PTG | Kan | — |
| — | pREDusk-*Ds*Red | *Ag*PSM*Dr*PTG | Kan | — |
| — | pREDusk-*Ds*Red | *Pa*PSM*Dr*PTG | Kan | — |
| — | pREDusk-*Ds*Red | *Rp*PSM*Dr*PTG | Kan | — |
| — | pREDusk-*Ds*Red | *Ag*PSM$_{-3}$ | Kan | — |
| — | pREDusk-*Ds*Red | *Pa*PSM$_{-10}$ | Kan | — |
| pFREDusk | pREDusk-*Ds*Red | *Pa*F1 | Kan | — |
| pFREDusk | pREDusk-*Ds*Red | *Pa*F1 | Strp | — |
| pFREDusk | pREDusk-*Ds*Red | *Pa*F1 | Amp | — |
| — | pREDusk-*Ds*Red | *Pa*F1$_{+34}$ | Kan | — |
| | pET21b(+) | *Dr*F1 | Amp | [8] |
| | pET21b(+) | *Dr*F1$_{-6b}$ | Amp | [9] |
| | pET21b(+) | *Ag*PSM | Amp | — |
| | pET21b(+) | *Pa*PSM | Amp | — |
| | pET21b(+) | *Rp*PSM | Amp | — |
| | pET21b(+) | *Ag*PTG | Amp | — |
| | pET21b(+) | *Pa*PTG | Amp | — |
| | pET21b(+) | *Rp*PTG | Amp | — |
| | pET21b(+) | *Ag*PSM-*Dr*PTG | Amp | — |
| | pET21b(+) | *Pa*PSM-*Dr*PTG | Amp | — |
| | pET21b(+) | *Rp*PSM-*Dr*PTG | Amp | — |
| | pET21b(+) | *Ag*NTS | Amp | — |
| | pET21b(+) | *Ag*NTS-*Ag*PTG | Amp | — |
| | pET21b(+) | *Pa*NTS | Amp | — |
| | pET21b(+) | *Pa*NTS-*Pa*PTG | Amp | — |
| | pET21b(+) | *Pa*F1 | Amp | — |
| | pET21b(+) | *Pa*F1$_{+34}$ | Amp | — |

***Table S4:*** ***Reverse primers used for PATCHY creation of PaPSM linker library.*** *Forward primers are listed in Meier et al.*[9].

| Primer | Sequence | Primer | Sequence |
|---|---|---|---|
| Pa_3742_rev | CTGCAGCGGATTACGC | Pa_3703_rev | CAGACGCTGACGCATAC |
| Pa_3739_rev | CAGCGGATTACGCAGATC | Pa_3700_rev | ACGCTGACGCATACGATC |
| Pa_3736_rev | CGGATTACGCAGATCATG | Pa_3697_rev | CTGACGCATACGATCAAC |
| Pa_3733_rev | ATTACGCAGATCATGACC | Pa_3694_rev | ACGCATACGATCAACTTCC |
| Pa_3730_rev | ACGCAGATCATGACCC | Pa_3691_rev | CATACGATCAACTTCCGC |
| Pa_3727_rev | CAGATCATGACCCAGAACTG | Pa_3688_rev | ACGATCAACTTCCGCC |
| Pa_3724_rev | ATCATGACCCAGAACTGC | Pa_3685_rev | ATCAACTTCCGCCGCATG |
| Pa_3721_rev | ATGACCCAGAACTGCAATC | Pa_3682_rev | AACTTCCGCCGCATGG |
| Pa_3718_rev | ACCCAGAACTGCAATCAG | Pa_3679_rev | TTCCGCCGCATGGTTG |
| Pa_3715_rev | CAGAACTGCAATCAGACG | Pa_3676_rev | CGCCGCATGGTTGAGG |
| Pa_3712_rev | AACTGCAATCAGACGC | Pa_3673_rev | CGCATGGTTGAGGCAC |
| Pa_3709_rev | TGCAATCAGACGCTGAC | Pa_3670_rev | ATGGTTGAGGCACAGTTC |
| Pa_3706_rev | AATCAGACGCTGACGC | Pa_3667_rev | GTTGAGGCACAGTTCCATC |
